# Investigating the activity of Varicella Zoster Virus (VZV) SUMO-targeted Ubiquitin Ligase ORF61

**DOI:** 10.64898/2026.03.11.710994

**Authors:** Anindita Puri, DSS Hembram, R Aravind, Ranabir Das

**Affiliations:** National Centre for Biological Sciences, Tata Institute of Fundamental Research Bengaluru, India

**Keywords:** Varicella Zoster virus (VZV), SUMO-Targeted Ubiquitin Ligase (STUbL), E3 Ligase, Protein Mimicry, ORF61

## Abstract

Varicella Zoster Virus (VZV) is a dsDNA virus that infects dermal cells and causes characteristic cutaneous lesions. The virus undergoes neurotropism and later causes secondary cycles of infection. In the host nucleus, Promyelocytic Leukaemia Nuclear Bodies (PML-NBs) spontaneously form around the VZV genome to repress viral gene expression. VZV encodes for a ubiquitin E3 ligase ORF61 to disperse PML-NBs and alleviate repression. ORF61 functions as a ubiquitin E3 ligase with a conserved RING domain at the N-terminal end. It carries three SUMO-interacting motifs (SIMs) that mediate interactions with SUMOylated proteins within PML bodies. The mechanism by which ORF61 disperses PML-NBs is poorly understood. To understand how ORF61 interacts with SUMOylated proteins, we investigated its interaction with SUMO and studied its SUMO-Targeted Ubiquitin Ligase (STUbL) activity. Our studies reveal that ORF61 co-opts the E2D family for ubiquitination activity. A specific network of interactions between the E2 enzyme, ORF61, and Ub facilitates polyubiquitination. ORF61 can synthesize branched polyubiquitin chains of K11, K48, and K63 linkages. The C-terminal SIM in ORF61 is a high-affinity binder of SUMO chains. Utilizing the SIM, ORF61 targets specific lysines on SUMO chains for ubiquitination. These studies provide crucial insights into the functional mechanism of viral STUbL ORF61.

## Introduction

Ubiquitination is a prevalent post-translational modification (PTM) that regulates multiple processes within the cell, such as protein turnover, signal transduction, DNA repair, cell cycle, and immune response [1]. It involves the covalent attachment of ubiquitin onto the substrate via an isopeptide bond, which is mediated by an enzymatic cascade comprising E1 (ubiquitin-activating enzymes), E2 (ubiquitin-conjugating enzymes), and E3 (ubiquitin ligases). There are 2 E1s, 32 E2s, and ∼670 E3s in humans. The E3 ligases regulate the substrate selectivity for ubiquitin modification and are classified into three families – RING (Really Interesting New Gene), HECT (Homologous to E6-AP Carboxyl Terminus), and RBR (RING Between RING) based on the different mechanisms of transferring ubiquitin on the substrate from E2 [2], [3]. Ubiquitin itself has seven lysines (K6, K11, K27, K29, K33, K48, and K63) and an N-terminal methionine, which can serve as acceptor sites. This gives rise to the formation of polyubiquitin chains of varying topologies, ranging from homotypic, heterotypic, and mixed to branched, which in turn regulate downstream signalling and direct substrates toward diverse functional outcomes. Deubiquitinases (DUbs) regulate the cascade by pruning the ubiquitin chains in a linkage-selective or promiscuous manner [2], [4].

Given the central role in cellular homeostasis, the ubiquitin system is a frequent target of many pathogens. Several viruses co-opt or mimic host ubiquitin ligases or deubiquitinases and integrate into the host signalling pathways to suppress antiviral responses or activate proviral responses. This strategy is particularly well studied for the human herpesviruses (HHVs) [5]. These enveloped double-stranded DNA viruses establish lifelong infections characterized by lytic replication cycles and latent neurotropic persistence. There are 8 HHVs categorised into three sub-families (α, β, and γ), and members of the α – HHVs are known to be quite prevalent in the human population. It includes Herpes Simplex Virus (HSV) types 1 and 2 and Varicella Zoster Virus (VZV). Their genomes enter the host nucleus during infection, where they encounter intrinsic host antiviral defences. One of the early host defences encountered by the viral genome is the viral DNA sensor IFI-16 (Interferon γ Inducible protein 16), which is recruited to the site of viral genome entry at the nuclear periphery. Another host immune response, the Promyelocytic Leukaemia Nuclear Bodies (PML-NBs), spontaneously form around the viral genome to restrict infection by sequestering and silencing the genome [6], [7]. To overcome this, HHVs have evolved to encode effector proteins that either prevent the formation of nuclear bodies or disperse them by nuclear body components via the ubiquitin-proteasome pathway [8], [9]. HSV encodes an immediate-early protein referred to as infected cell protein 0 (ICP0), which functions as a special class of RING E3 ligase called a SUMO-targeted ubiquitin ligase (STUbL). Proteins modified by another ubiquitin–like protein, Small Ubiquitin–like Modifier (SUMO), are ubiquitinated by STUbLs. These ligases include a RING domain for ligase activity and a SUMO-interacting motif (SIM) to specifically interact with SUMOylated proteins. HSV-encoded ICP0 protein induces ubiquitination and subsequent proteasomal degradation of SUMOylated PML and Sp100, thereby dispersing PML-NBs, enabling viral gene expression, and initiating pathogenesis [10], [11]. ICP0 has also been reported to degrade the IFI-16 filament complex, which traps viral DNA to prevent entry into the nucleus [7], [12].

The mechanisms by which other members of the alphaherpesvirus family engage the ubiquitin–SUMO axis to counteract PML nuclear bodies are poorly understood. Within the alphaherpesviruses, the STUbL ICP0 expressed by HSV-1 and HSV-2 is conserved with approximately 60% sequence identity. The ortholog ORF61 expressed by VZV contains an N-terminal RING domain and functions as an E3 ligase similar to ICP0. However, the sequence conservation between ICP0 and ORF61 is merely 10%, suggesting a divergence in their functional mechanisms. While ICP0 includes one SIM, ORF61 contains three SIMs. While ICP0 degrades PML-NBs via the ubiquitin-proteasome pathway, ORF61 disperses PML-NBs through an unknown mechanism. In this study, we examined the interaction and ubiquitination activity of ORF61 with host E2 enzymes. Furthermore, we investigated its interaction and STUbL activity with the substrate SUMO. These studies indicate that ORF61 can form both linear and branched ubiquitin chains on substrate SUMO chains. The ubiquitination target sites on SUMO are specific and resemble those of host STUbLs, suggesting that ORF61’s activity mimics that of host STUbLs. Finally, we propose ORF61 targets upstream factors like IFI-16 for degradation to prevent de novo PML-NB formation.

## Materials and Methods

### Plasmids

Full-length ORF61 sequence was synthesised from Invitrogen and sub-cloned in pET15b expression vector with BamH1and Nde1 restriction sites. The RING and SIM mutants were generated by site-directed mutagenesis using primers synthesised by Eurofins. Constructs for tetra-SUMO2 and hs-RNF4 were kindly provided by Dr. Cynthia Wolberger.

### Peptides

For NMR titration studies, individual ORF61 SIM peptides were synthesised and purchased in lyophilised powder from Lifetein LLC.

### Protein expression and purification

The ORF61 construct was transformed and expressed in the *E. coli* BL 21 strain. The cells were grown at 37°C to an OD600 of 0.8 and induced with 1mM IPTG for 4 hours. The cells were harvested and stored in -80°C until processed. The harvested cells were lysed in 50 mM HEPES (pH 7.5), 300 mM NaCl, 2% glycerol, 2 mM DTT, 0.1% Tween 20 with one anti-protease tablet (Roche). Cells were sonicated at 40% amplitude for 10 seconds. The lysate was clarified at high-speed centrifugation at 14,000 RPM at 4°C. The pellet obtained was resuspended in 50 mM HEPES (pH 7.5), 300 mM NaCl, and 6 M Urea, and left overnight at 4°C on a shaker. The resuspension mix was sonicated the following day for 5 minutes and clarified by high-speed centrifugation at 14,000 RPM for 20 minutes at 4 °C. The supernatant obtained was incubated with Ni-NTA beads equilibrated in 50 mM HEPES (pH 7.5), 300 mM NaCl, 6 M Urea buffer for one hour at 4°C. This was followed by extensive washes with 50 mM HEPES (pH 7.5) and 300 mM NaCl, and a gradient batch elution with increasing imidazole concentrations up to 500 mM. It enabled partial on-column refolding via buffer exchange. The eluted fractions containing the protein of interest were pooled and diluted to a higher volume with 50 mM HEPES (pH 7.5), 300 mM NaCl, 2% glycerol, and 100 mM L-Arginine buffer, thereby reducing the urea concentration to 300 to 500 mM. The remainder of the buffer exchange to remove urea and subsequently arginine was carried out by centrifugal filtration using Amicon concentrators, with the final buffer being 50 mM HEPES (pH 7.5) and 300 mM NaCl. The protein was then aliquoted and stored at -80°C by flash-freezing.

E1, E2, Ubiquitin, ^15^N SUMO1 and ^15^N SUMO2 were expressed and purified as discussed in previous studies [11]. Ubiquitin S20C was purified in the same way as ubiquitin. RNF4 and tetra-SUMO2 were purified as previously reported [13].

### Fluorescence tagging of substrate

For polyubiquitination and the STUbL assay, ubiquitin S20C and tetra-SUMO2 (the respective substrates) were tagged with Alexa Fluor 488 Maleimide (Invitrogen). The substrate was incubated with the dye at a molar ratio based on the protein concentration at 4°C for 2 hours. The excess dye was removed via dialysis, followed by passing the labelled protein through a desalting column (PD MiniTrap G-25 - cytiva). SDS-PAGE was resolved to evaluate for any excess untagged fluorophore.

### In vitro Biochemical Assays

The *in vitro* polyubiquitination assay was conducted using 1.5 µM E1, 4 µM E2, 15 µM E3 (Purified ORF61), and 30 µM Alexa Fluor Maleimide (Invitrogen)-tagged Ubiquitin S20C in the presence of 2 mM ATP in reaction buffer containing 50 mM Tris (pH 8.5), 50 mM NaCl, 2 mM MgCl2, 0.4 mM DTT and 5 mM ATP. The reaction was incubated at 37°C. Samples were taken at different time points and mixed with a non-reducing 5x non-reducing SDS-PAGE loading dye. The samples were resolved in 12% SDS-PAGE gel. Fluorescence signal readout was visualised using an Amersham Typhoon instrument.

The *in vitro* SUMO Targeted Ubiquitin Ligase (STUbL) assay was carried out in the presence of 1.5 µM E1, 4 µM E2, 15 µM E3, 100 µM Ubiquitin, 5 µM Alexa Fluor Maleimide (Invitrogen)-tagged TetraSUMO2 in the presence of 2mM ATP in the reaction buffer mentioned. The reaction was incubated at 37°C for up to 3 hours. Samples were taken at different time points and mixed with a non-reducing 5x non-reducing SDS-PAGE loading dye and resolved on a 12% SDS-PAGE gel. Gel was visualised using fluorescence on an Amersham Typhoon instrument. Fluorescent band intensities were measured using ImageJ [14] and normalised with respect to the no E1 lane (negative control) to calculate the fold change differences among the wild type and mutants. 2-tailed, equal variance Student T-test was performed to calculate the significance of measured changes.

### E2 Screening

E2 screening was performed *in vitro* using the commercial screen kit supplied by RnD Systems - E2Select Ubiquitin Conjugation Kit (K-982) manufactured by Boston Biochem, Inc. The assay was conducted using ORF61 as the E3 ligase of interest and Alexa Fluor 488-tagged Ubiquitin. The screening was set up with no E3 as a control and incubated at 37°C for 30 minutes. Samples were mixed with non-reducing 5x non-reducing SDS-PAGE loading dye and resolved in 12% SDS-PAGE gel. Fluorescence signal readout was visualised using an Amersham Typhoon instrument.

### NMR Experiments

NMR spectra were acquired at 298K on an 800 MHz Bruker Avance III HD spectrometer with a cryoprobe head, processed with NMR pipe [15], and analysed with NMRFAM-SPARKY [16]. A sample was prepared with labelled SUMO (150-250 µM) supplemented with 10% D2O, and an increasing volume of peptide (∼3000 to 7300 µM stock) was added as per increasing ratios. 15N-edited Heteronuclear Single Quantum Coherence (HSQC) experiments were recorded at each ratio. From the chemical shifts recorded at each ratio, the binding dissociation constant was calculated based on the 1:1::Protein:Ligand model using the equation CSP_obs_ = CSP_max_ {([P]t + [L]t + Kd) - [([P]t + [L]t + K_d_)^2^-4[P]t[L]t]^1/2^}/2[P]t, where [P]t and [L]t are total concentrations of protein and ligand at any titration point.

### Modelling E2D2-ORF61-Ub complex

The sequence of the RING domain in ORF61 (GSS..PSDDD) was obtained from the UniProt database [17], and its structure was predicted using AlphaFold [18]. Two zinc-coordination complexes were subsequently modelled on the ORF61 structure using the MIB2 server [19], [20] which identifies, and places metal coordination sites based on structural templates. To construct the E2D2–ORF61–ubiquitin complex, the crystal structure of E2D2 in complex with the RING domain of RNF38 and ubiquitin in the closed conformation (PDB ID: 4V3K) [21] was used as a template. ORF61 was structurally aligned onto the RING domain of RNF38, and the latter was then removed to generate the final E2D2–ORF61–ubiquitin complex using UCSF Chimera [22]. The resulting complex structure was processed and cleaned using pdb4amber to prepare it for molecular dynamics simulations.

#### System Preparation

All molecular dynamics (MD) simulations were performed using the AMBER18 [23] software suite with the ff14SB force field [24] for protein atoms and the TIP3P [25]. An explicit water model for solvation. Parameters for zinc coordination were incorporated from a customized version of the Zinc AMBER Force Field (ZAFF) [26], while thioester connectivity was modelled using previously developed modified parameters. Coordinate bonds were manually defined to reflect the tetrahedral geometry of zinc binding, involving cysteine sulphur atoms and, in one site, a histidine imidazole nitrogen. Nonphysical bonds, including unintended Zn–Zn interactions, were explicitly removed. The system was solvated in a cubic box of TIP3P water molecules with a 12 Å buffer around the solute. Sodium ions were added to neutralise the system’s net charge. To enable stable simulations with a 4 fs integration timestep, hydrogen mass repartitioning (HMR) [27] was applied.

#### System Equilibration

Energy minimisation was carried out in two stages. In the first stage, only solvent molecules and counterions were relaxed while the protein was harmonically restrained (20 kcal/mol/Å²). This was followed by a second minimisation step in which all atoms in the system were allowed to move freely. Both minimisation steps used steepest-descent followed by conjugate-gradient algorithms until convergence. Subsequent heating and equilibration were carried out through a staged protocol. The system was gradually heated from 0 K to 300 K over 100 ps under the NVT ensemble, with harmonic restraints (20 kcal/mol/Å²) applied to all protein heavy atoms. Temperature regulation was achieved using Langevin dynamics with a collision frequency of 5 ps⁻¹. This was followed by 100 ps of equilibration under NPT conditions (1 atm, 300 K), with the same restraint scheme. The pressure was controlled using the Berendsen barostat with a relaxation time of 1.0 ps. To facilitate conformational relaxation, the system was subjected to three short MD phases in which the restraint strength on protein backbone atoms was sequentially reduced from 10 to 1 kcal/mol/Å². These simulations were carried out with a timestep of 2–4 fs using SHAKE constraints on bonds involving hydrogen atoms. The final equilibration phase was run without restraints for 400 ps under NPT conditions to ensure complete relaxation of solvent and side chains.

#### Production

Production MD simulations were performed in five independent replicas, each initiated from the final equilibrated structure with randomised initial velocities. Each replica was simulated for 1 μs, yielding an aggregate sampling time of 5 μs. Simulations were carried out in the NPT ensemble at 300 K and 1 atm. Temperature coupling was maintained using Langevin dynamics, and pressure coupling was applied using isotropic scaling via the Berendsen barostat. A 4 fs integration timestep was employed in all production runs. Long-range electrostatics were treated using particle mesh Ewald (PME) with a real-space cutoff of 10 Å. SHAKE constraints were applied to all bonds involving hydrogen atoms, and periodic boundary conditions were imposed with coordinate wrapping enabled. Trajectory frames were recorded every 20 ps, giving 50000 frames per replica.

#### Analysis

Following production simulations, the trajectories from each of the five independent replicas were post-processed using cpptraj [28]. Frames were extracted from each trajectory at intervals corresponding to every fifth frame, starting from the first to the final snapshot. All water molecules and counterions were removed, and periodic boundary conditions were resolved using the “autoimage” function to centre and reimage the solute. The coordinates were aligned to the initial frame based on backbone heavy atoms. The resulting solvent-free trajectories were saved in a combined file format suitable for downstream analyses.

#### Contact Maps

Intermolecular residue–residue contacts were computed using the native contacts module in cpptraj, considering all-atom contacts formed between pairs of residues within 7 Å. Specifically, two distinct interaction interfaces were analysed: (i) between the RING domain of ORF61 and ubiquitin, and (ii) between the RING domain of ORF61 and E2D2. For each pair, both native and non-native contacts were computed, yielding four datasets, one representing native and the other non-native maps for each interaction.

To obtain a comprehensive view of the contact network, native and non-native contact frequencies were merged by summing the corresponding contact probabilities for each residue pair. The resulting contact values were normalised by the maximum contact frequency observed in each interaction map, yielding values in the range of 0 to 1. Normalised contact data were visualised as 2D heatmaps using custom Python scripts.

#### Clustering & Interaction Analysis

To identify representative conformations from the simulation ensemble, hierarchical agglomerative clustering [29] was performed using cpptraj. Clustering was based on RMSD of backbone and side-chain heavy atoms (C, N, O, CA, CB). The clustering protocol employed average-linkage as the linkage criterion, with an epsilon value of 3.0 Å and a maximum of 10 clusters. A frame sieve interval of 10 was applied to reduce computational cost, with frames randomly selected for clustering. For each cluster, an average structure and a representative structure (centroid) were extracted. The representative structure from the first cluster, which contained the largest number of frames and exhibited the lowest average distance to the centroid (1.73 Å), was selected for further analysis. This representative structure was subjected to a short energy minimisation protocol to relieve any potential steric clashes prior to downstream analysis. The minimised structure was then used as input for residue-level interaction analysis with LigPlot+ [30] allowing visualisation of key protein–protein and protein–ligand contacts in the most probable conformational state of the complex.

### Mass Spectrometry based Investigation and Analysis

The *in vitro* polyubiquitination assay was conducted using 1.5 µM E1, 4 µM E2, 15 µM E3 (purified ORF61), and 30 µM ubiquitin in the presence of 2 mM ATP in reaction buffer. The reaction was incubated at 37°C for 1 hour. The samples were then incubated with purified clippase enzyme for 2 hours at 37°C.

For linkage detection: The samples were quenched after incubation by addition of reducing SDS-PAGE sample dye and resolved in 12% SDS-PAGE gel. The mono-ubiquitin band was visualised and cut after Coomassie brilliant blue R-250 staining and destaining. The band was chopped into smaller pieces and destained extensively in 1:1::TAEB:ACN buffer followed by dehydration in 100%ACN. The dehydrated gel pieces were subjected for overnight trypsin digestion (1:20 based on protein concentration) at 37°C. The peptides were eluted in 1:0.1::ACN:FA mix and vacuum dried. The peptides were reconstituted in 0.1%FA and quantified prior injection. The same steps were followed for detection of lysines modified in the STUbL assay.

For chain architecture: 5µL of ubiquitination assay samples were injected in Agilent Zorbax C8 analytical HPLC column (Zorbax-C8883995-906) for intact mass assessment in LTQ Orbitrap XL with API source and a run time of 25 minutes. The peaks obtained were assessed for masses corresponding to ubiquitin and ubiquitin carrying one or multiple di-glycine (-GG) motif. Based on the charged state observed, the mass was back calculated for the species visible in the spectra.

LC-MS/MS analysis followed established parameters [31]. Briefly, 1 µg samples were injected into Easy nLC-II and separated on an Agilent ZORBAX 300SB-C18 (15 cm, Cat. No.5065-9911) using a 60-min step-gradient of 0.1% formic acid and 90% acetonitrile, starting from 10% B to 35% B in 53 mins. Data were acquired on an Orbitrap Fusion in DDA mode: MS1 resolution 120K (AGC 4e5, max IIT 50 ms, 375–1200 m/z) profile mode. MS2 was acquired using IonTrap rapid mode (AGC 1e4, IIT 30 ms, NCE: HCD30) using quadrupole isolation window of 1.6m/z with parallelization of ion accumulation. The Duty cycle time set at 3sec and dynamic exclusion was set for 30sec. Data was searched using Proteome Discpoverer 2.2 using Sequest HT search algorithm against Human Swissprot database (2025_07_30). Trypsin digestion parameters we set at 2 missed cleavage, peptide length 6–144 AA, with 10ppm precursor and 0.6 Da mass tolerance. Cysteine-Carbamidomethylation (fixed) and methionine-oxidation (variable), N & Q-Deamidation (variable) and Lysine - Ubiquitination (Variable) were applied. Peptide and Protein and PTM FDR was controlled at 1% using Percolator, Protein FDR validator and PhosphoRS Nodes. Precursor Ion Quantifier node was used for peptide and protein label free quantitation based on peptide intensities.

## Results

### ORF61 co-opts the E2D family of host E2 Enzymes for ubiquitination

We validated the ubiquitin RING E3 ligase activity for the recombinant ORF61 through an *in vitro* ubiquitination assay, conducted using human ubiquitin-activating enzyme UbE1 (E1), ubiquitin-conjugating enzyme (E2) Ube2d2/Ubch5b, purified ORF61 as the E3, and fluorescently labelled Ub. After the reaction, the samples were resolved on an SDS-PAGE gel, and the fluorescent polyubiquitin (polyUb) bands were imaged. Within the first few seconds (0 min lane), a diubiquitin band emerged, which processively led to higher-molecular-weight polyUb bands within 30 min of reaction (Fig 1a and Fig 1Sa), confirming ORF61’s robust E3 ligase activity.

**Figure 1:**
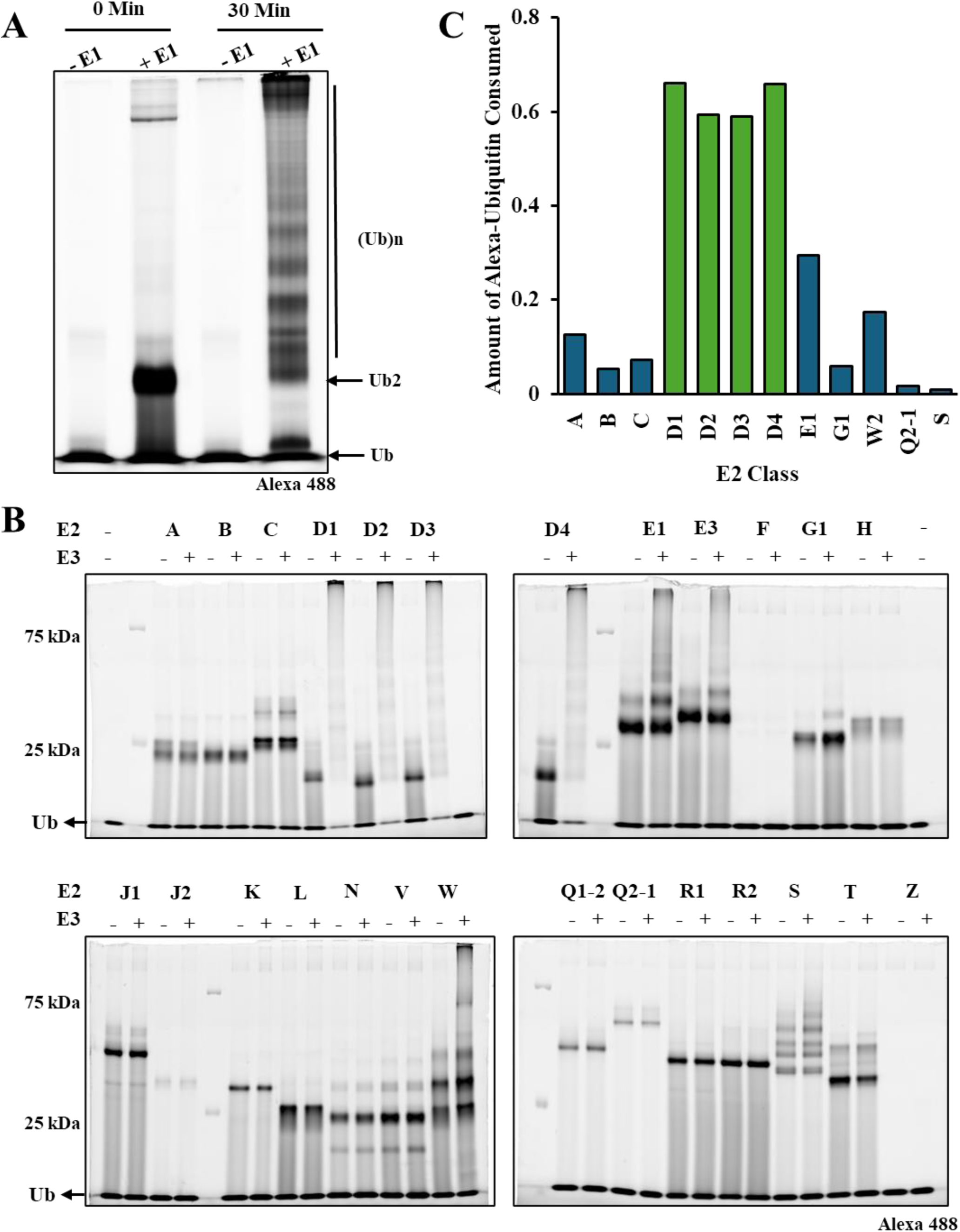
Investigating activity of ORF61 with host E2 Enzymes. A) Polyubiquitination activity of purified recombinant ORF61. The assay was conducted using Alexa – 488 labelled ub at 37°C in presence of E1 and E2D2. Reaction without E1 (No E1) was a negative control. The emergence of polyubiquitin bands over 30 minutes was visualized through fluorescence imaging and suggested active RING E3 ubiquitin ligase activity. B) 26 E2 enzymes were screened with the purified recombinant ORF61 to evaluate for the potential E2 enzymes that can function with it. The assay readout at 30 minutes visualized using fluorescence since Alexa – ub was used. C) Plot shows the amount of Alexa – ubiquitin consumed in each reaction with select E2 enzymes by comparing the ratio of ubiquitin band intensity for with and without E3 lanes for the E2.

Subsequently, an *in vitro* E2 screening was performed to assess for all possible human E2-enzymes that could be potentially co-opted by ORF61 for ubiquitination. The above ubiquitination assay was repeated in the screen with 26 known human E2 enzymes. ORF61 formed ubiquitin chains robustly with the E2D family, while faint bands at higher molecular weights were observed with the E2E1, E2E3, and E2W2 families (Fig 1b and 1Sb). From ratio metric analysis, close to 60% of free monoubiquitin was consumed in the case of the E2D family, while only ∼30% for E2E1 and ∼20% for E2W2 (Fig 1c and Fig 1Sc). The E2E family can function independent of E3 [32], which can be observed by the presence of polyUb bands in the no E3 (-E3) lane. The presence of ORF61 appears to subtly enhance polyUb synthesis in E2E, a phenomenon also observed in E2W. Overall, ORF61 functions robustly with the E2D family for polyubiquitination, and these host E2 enzymes may be co-opted by ORF61 during VZV infection.

### The ORF61-RING:E2∼Ub Complex

To understand how ORF61 functions with the E2D enzymes, we studied its interaction with the E2∼Ub conjugate by modelling. To this end, we chose the well-studied E2 enzyme E2D2 as a representative of the E2D family. The structure of ORF61 is unknown. AlphaFold predicted that ORF61 is predominantly disordered, with only the RING domain and the central hydrophobic stretch predicted with confidence. The predicted ORF61-RING domain has a conserved core of a ββα (or β2α) fold with C3HC4 cross-braced zinc-ion coordination scheme (Fig. 2Sa). The structure is similar to that determined for Equine Herpes virus which is also known to resemble the Herpes Simplex virus type 1 protein Vmw110 (ICP0) [33], [34]. It is interesting to note that despite poor sequence conservation between ICP0-RING and ORF61-RING, their structures are conserved. Using the ORF61-RING predicted structure, we modelled the ORF61-RING:E2D2∼Ub complex structure. The solved crystal structure of the RNF38:E2D2∼Ub complex (PDB-ID 4V3K) was used as a template, with the RNF38 RING substituted by the ORF61 RING domain. The structure was further stabilized and refined by MD simulations to obtain the final ORF61-RING:E2D2∼Ub complex (Fig 2b).

**Figure 2:**
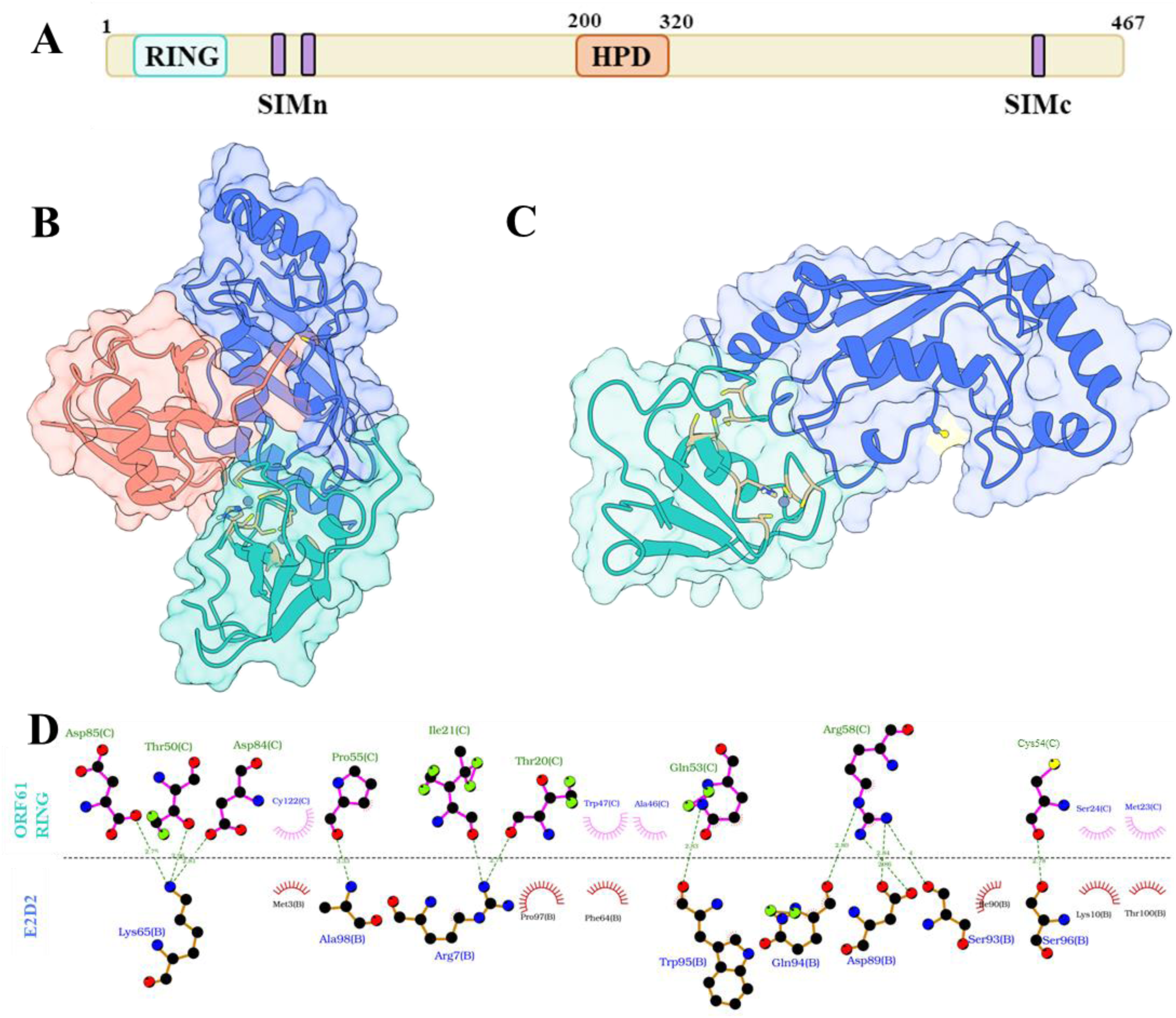
Structure Modelling and Interface Analysis for ORF61 RING:E2∼Ub Complex. A) Domain organization of ORF61 wherein RING domain with classic ββ⍺ fold and cross-braced zinc coordination scheme is present at the N – terminal. The three SUMO Interacting Motifs (SIMs) are distributed across ORF61 such that two are near the RING domain and one is at the C– terminal end of the protein. There is hydrophobic domain (HPD) at the center of the protein which is likely to aid in self-oligomerization. B) Model of ORF61 RING Domain:E2D2∼Ub complex is oriented to highlight the conserved mode of interaction. The ORF61 RING domain is coloured in sea green, E2D2 in blue and ubiquitin in light pink. C) The model is presented without ubiquitin to highlight the canonical interaction interface between the ORF61 RING domain and E2D2. The RING domain has the zinc ions shown in dark grey and the cysteine and histidine residues coordinating to the zinc ions are coloured in tan. The active site cystine in E2D2 is coloured in yellow. D) The interaction map shows the residues of ORF61 RING and E2D2 at the interface interacting through hydrophobic (semi-circle representation) and hydrogen bond (dotted line representation) interaction. The atoms are colour coded such that the solid black circles are carbon, red are oxygen, blue are nitrogen and green are sidechain hydrogen atoms. The Sulphur in the cystine residue involved in zinc coordination is coloured in yellow.

The structure showed extensive interface between ORF61-RING and E2D2, where loops 4 and 7 and helix 1 of E2D2 interact with two loop regions in between zinc-coordinating residues of the RING Domain [35], [36]. The complex has the E2∼Ub in the closed state (Fig 2c). Based on contact maps from simulation studies, a detailed network of residue interactions was reconstructed for the complex (Fig 2d and Fig. 2S). The RING domains have a strict sequence conservation of the zinc-coordinating cystine and histidine residues [35]. A conserved hydrophobic residue (I/ L or V) is typically present within the first two cysteines in the RING domain, which forms a critical interaction with the E2 enzyme. The corresponding residue is I21 in the ORF61 RING domain, which interacts with the R7 sidechain in helix 1 of E2. A key linchpin [LP] arginine, present immediately after the last zinc-coordinating cysteine, allosterically brings the E2∼Ub conjugate into a closed state for efficient ubiquitin transfer [37]. In the ORF61 RING domain, R58 was identified as the LP, and it interacted with the backbone carbonyls of E2 residues involving Q94, D89, and S93, as well as I36 in Ub. W47 of the ORF61-RING domain interacts with P97 in E2 via stacked conformation and with F64 via pi-pi interaction. Even though the individual residues of the RING domain differ, their interactions with E2D2 are similar to other RING:E2D complexes, such as RNF4:UbcH5a (PDB: 4AP4) and RNF38:UbcH5b (PDB: 4V3K), where the RING E3s are host ubiquitin ligases. This highlights the ease with which ORF61 integrates into the host ubiquitination pathway, as it can mimic and function similar to the host ubiquitin E3 ligase.

### The ORF61-RING:E2∼Ub contacts are critical for polyubiquitination activity

To study the relevance of contacts at the ORF61:E2D2∼Ub interface for ubiquitination, we selected four residues – I21, W47, L56, and R58 of the RING domain for targeted mutational study (Fig 3a). I21, W47, and R58 have direct contacts with E2 residues at the interface. L56 forms hydrophobic contacts with L8 in Ub. The selected residues were individually mutated to alanine via site-directed mutagenesis to disrupt their interactions with E2. Each mutant was then assessed for ubiquitin E3 ligase activity using a polyubiquitination assay and compared with wild-type ORF61. Reaction without E3 was used as a negative control to account for any basal E3-independent activity of E2D2 (Fig 3b and 3c, Fig 3S). The general trend observed across all mutants was reduced activity, likely due to weakened E2:E3 interactions and reduced activity of the RING:E2∼Ub complex. We observed that alanine mutants of the interface residues I21, W47, and R58 reduced activity by nearly 85%. Perturbing L56:L8 interaction reduced the activity by ∼50%. Therefore, the loss of RING:E2 and RING:Ub contacts perturb the activity of ORF61-RING:E2∼Ub complex, emphasising the importance of the specific interaction network at the interface.

**Figure 03:**
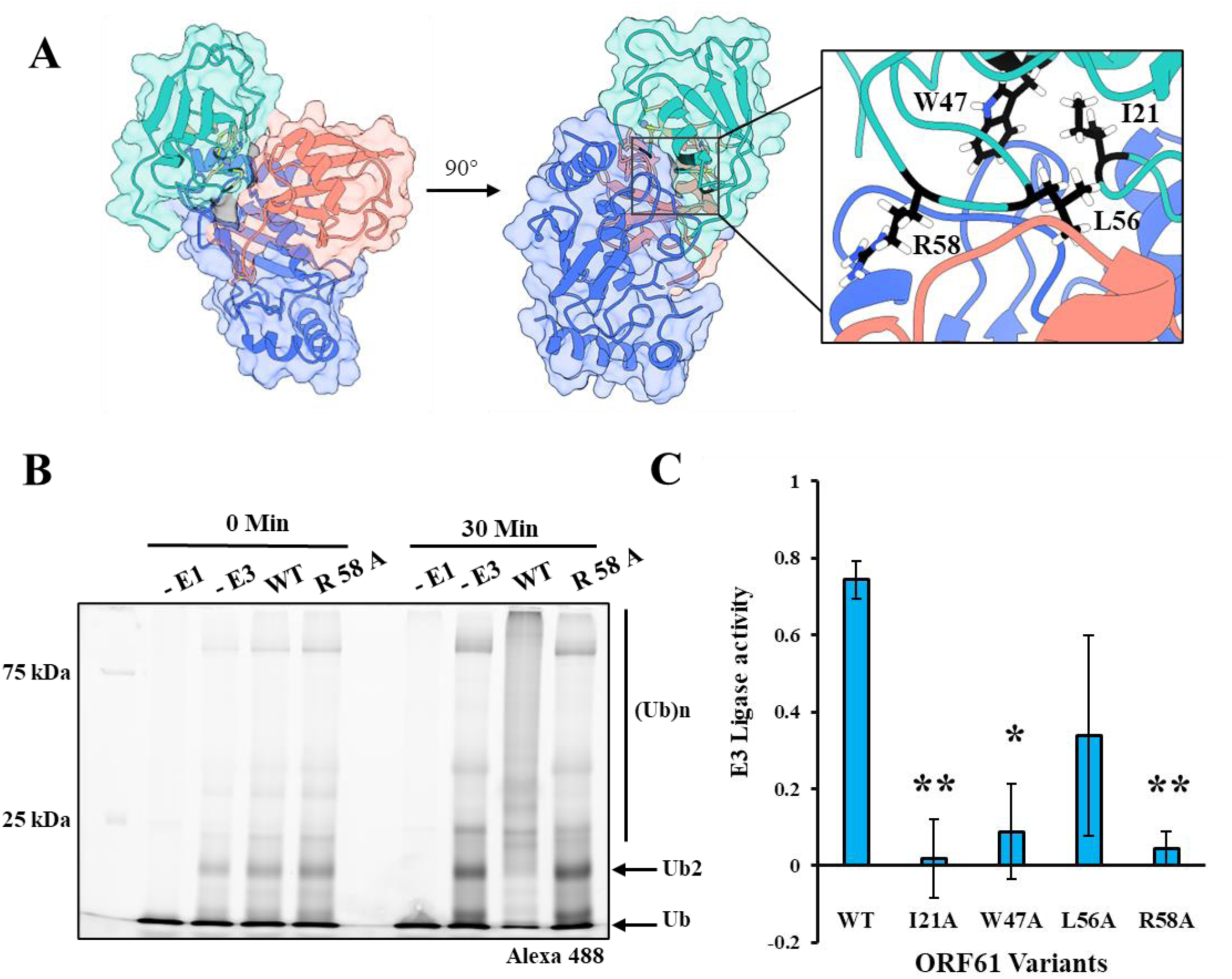
Assessment of Functional Relevance of Residues at the RING:E2 Interface. A) Surface visualization of 4 residues – I21, W47, L56 and R58 coloured in black on ORF61 RING at the RING (sea green) – E2D2 (blue) – Ubiquitin (light pink) complex. The residues were selected based on the assessment of residues involved at interface for other RING:E2 pairs and the type of interaction the residues engage in. The inset shows a zoomed in visualization of residues selected. B) Polyubiquitination activity for one of the RING mutants R58A compared to wild type ORF61. No E1 and No E3 were used as negative control. This helps to compare the extent of loss in E3 ligase activity of the mutant while comparing the basal E3 – independent activity of E2D2 in the assay. C) The plot shows the ORF61 E3 ligase activity based on ubiquitin consumed wherein the amount of ubiquitin consumed for each mutant has been plotted against the wild-type for comparison. The assays were done in duplicates.

### ORF61 can form branched ubiquitin chains

Polyubiquitin chains are of diverse topology ranging from linear, homotypic to heterotypic and branched with mixed linkages expanding the downstream signalling outcome. Pathogens often exploit this code. For example, NleL (Non-LEE encoded effector ligase) encoded by the enterohaemorrhagic *E. coli* (EHEC O157:H7) is injected into host cells to restrict number of actin-rich pedestal formed on cells under the adherent bacteria. NleL forms atypical branched K6 – K48 chains, which are implicated for promoting proteasomal degradation and inhibiting mitophagy [38], [39]. We were interested in the chain architecture formed by ORF61. We carried out ubiquitination assay with ORF61 as the E3, incubated the reaction with a clippase enzyme [40]. Clippases form a distinct class of non-traditional deubiquitinating enzymes wherein upon cleavage of ubiquitin and ubiquitin chains, a di-glycine motif (-GG) peptide is left on the substrate as the enzyme cleaves between the LR | GG making it possible to trace the site of ubiquitin modification on the substrate. This method is referred to as Ub-Clipping [40], [41].

Utilising this technique, we resolved the samples post treatment with clippase on SDS-PAGE gel and isolated the mono-ubiquitin band. The band was digested with trypsin. The peptides obtained were then detected using mass spectrometry. From the results, we observed that ORF61 catalysed the formation of K63, K48, and K11 linkages as the -GG remnant peptide was conjugated to the corresponding lysines (Fig 4a). Analysis of the monoubiquitin without trypsin digestion by intact mass analysis of the reaction after incubation with clippase detected monoubiquitin carrying two -GG remnant peptides, suggesting the presence of branched chains. About 52% of ubiquitin was either free, monoubiquitnated, or terminal ubiquitin at the chain end. About ∼32% was modified with single -GG indicating linear chain, and ∼ 16% corresponded to branched chains (Fig 4b and 4c). NLeL forms about 10-14% branched ubiquitin chains [42], which is comparable to ORF61, suggesting microbial effectors show similar trends in branching. Since ORF61 prefers to synthesized K11, K48 and K63 linkages, the ubiquitin branching could be either K48-K63, K11-K48, or K11-K63 chains, whose functional relevance is a subject of future research.

**Figure 4:**
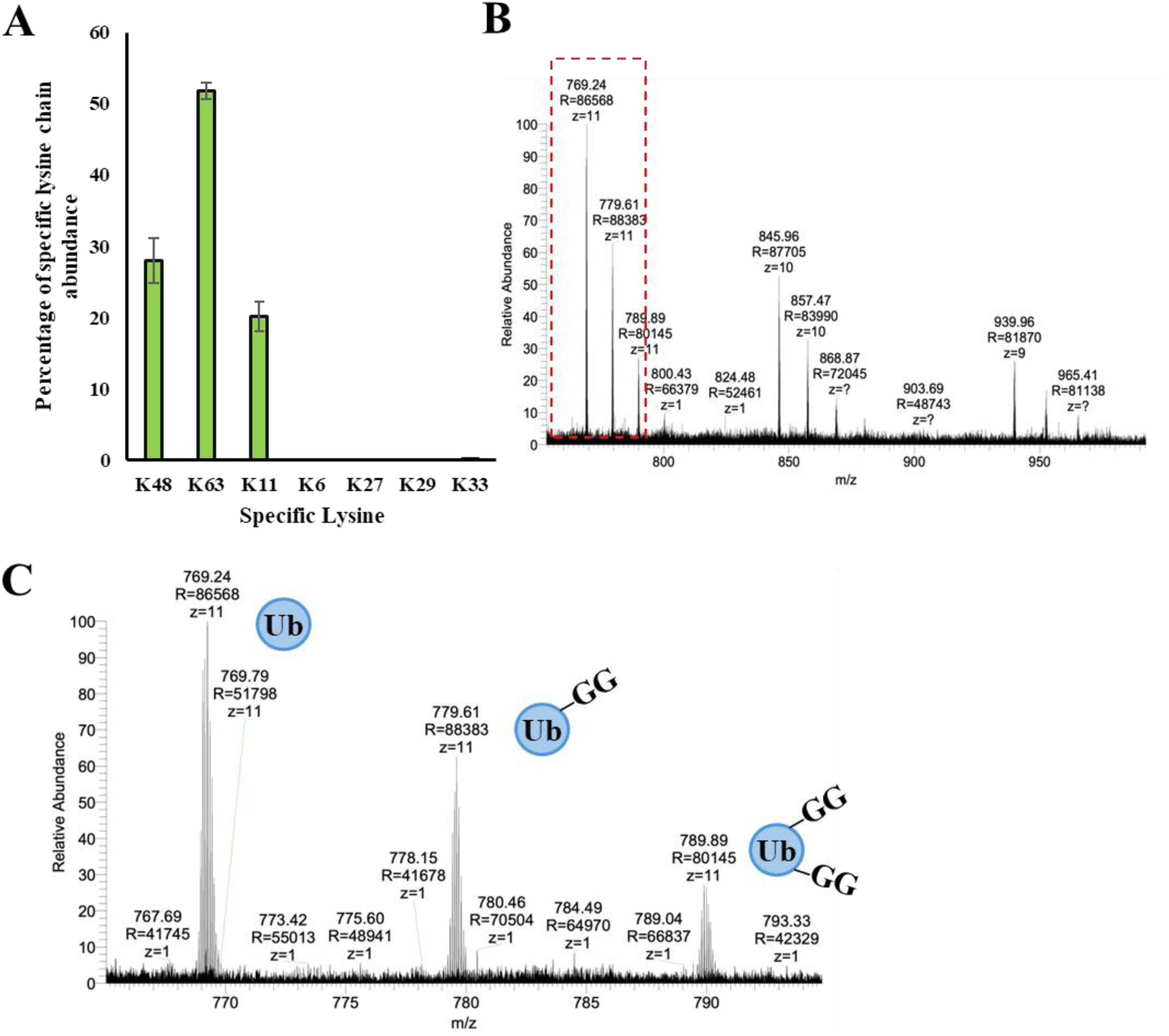
Investigating the lysine specificity and prevalence of branched architecture for ORF61. A) Polyubiquitin reaction was saturated and then subjected to ub-clipping using clippase enzyme. The monoubiquitin species were purified and processed for trypsin digestion. The peptides revealed the lysine with di-glycine remnant resistant to trypsin digestion. The peptides detected for specific linkage type are plotted here. B) The same polyubiquitin reaction was subjected to intact mass and we observed a predominant charged state. The three species with unmodified ubiquitin with highest relative abundance and about 30% carrying two di-glycine modifications suggestive of presence of branch point in the polyubiquitin species formed are shown in a zoomed inset highlighted in red in C).

### C-terminal ORF61 SIM Interacts with High Affinity with both SUMO Isoforms

Apart from the N-terminal RING domain, there are three SUMO-Interacting Motifs (SIMs) present in the disordered region of ORF61. Two of them are present near the RING domain – SIM N1 and SIM N2 (jointly referred to as SIMn), and the third SIM at the C-terminal end (SIMc) (Fig 2a). ORF61 localises to the nucleus through its C-terminal NLS [43]. The SIMs drive colocalization of ORF61 with PML-NBs, presumably via interaction with SUMOylated PML and other SUMOylated proteins in the PML-NBs [44]. To study the interaction of SIMs with SUMO1 and SUMO2 isoforms, we used NMR titration experiments.

Peptides of each SIM were titrated against ^15^N-labeled SUMO1 and SUMO2. The backbone chemical shifts of SUMO with increasing SIM peptide ratios on SUMO isoforms were studied using Nuclear Magnetic Resonance (NMR) and detected via ^15^N-edited Heteronuclear Single Quantum Coherence (HSQC) experiments. SIM binding alters the chemical environment of SUMO residues, as reflected in changes in the chemical shifts of backbone NH resonances (Fig. 5a). These chemical shift differences identify the region between β2 and α1 as the primary interface of SUMO:SIM interactions (Fig 5b-e). The titrations were repeated with N-terminal SIMs SIM N1 and SIM N2, which suggested a similar binding interface across all ORF61 SIMs (Fig. 5S).

**Figure 5:**
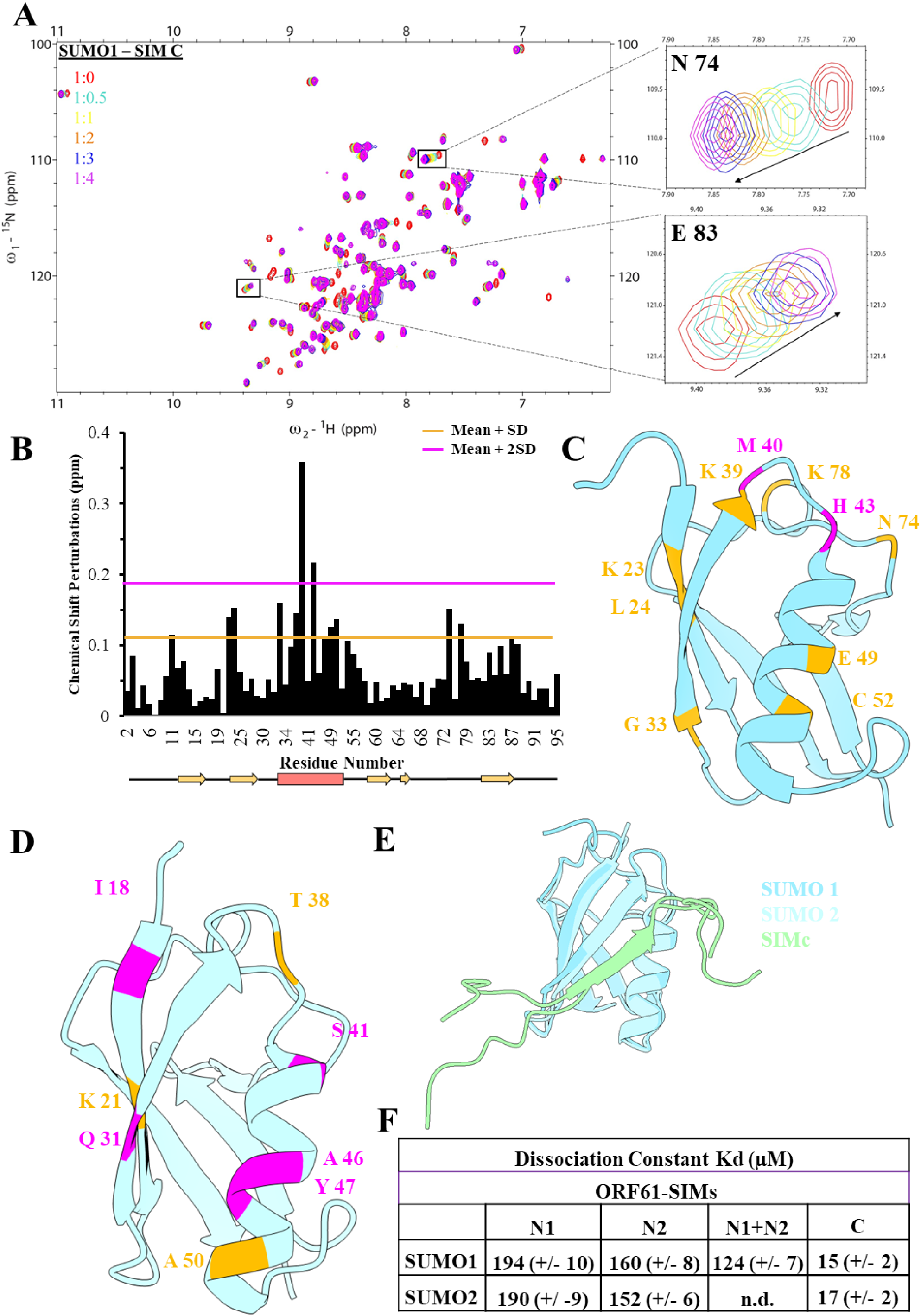
Validating interaction of ORF61 SIMs with SUMO isoforms. A) ^15^N-HSQC plot of SUMO1 isoform against increasing concentrations of ORF61 SUMO Interacting Motif (SIM) of C-terminal (SIMc) peptide at increasing concentrations. The inset shows two residues showing major perturbations. B) The chemical shift perturbation for all residues has been plotted. C) Surface representation of SUMO1 (2N1V) oriented at the β2-⍺ groove, also known as the hydrophobic groove of SUMO, has residues coloured differently based on the significant perturbations observed. D) The surface representation of SUMO2 (2N1W), also oriented at the hydrophobic groove, has residues coloured differently based on the significant perturbations observed in the titration of SUMO2 isoform against SIMc peptide. E) The AlphaFold predicted structure of the SIMc peptide interacting with the two SUMO isoforms superimposed where SUMO1 (light sky blue) and SUMO2 (light blue). The SIMc peptide (light green) adopts a parallel beta-strand conformation that sits at the hydrophobic groove of SUMO. F) The table shows the dissociation constant calculated for the two isoforms against the three SIM regions present in ORF61.

SUMO:SIM interactions are typically of the order of 100 μM [45], [46]. The change in chemical shifts with SIM concentrations was fitted to yield the binding dissociation constants (K_d_) for each ORF61 SIM (Fig 5f). SIM-N1 and SIM-N2 bind to SUMO1/2 with a K_d_ of 190 μM and 160 μM, respectively. However, the SIMc binds to SUMO1/2 with a K_d_ of 15 μM, suggesting a strong affinity. Notably, the SIMc motif is *LTIDL* and is a variant of a less frequent SIM motif [47]. This motif is unique for VZV ORF61, and not reported for any of the other ICP0 orthologs among the other alpha herpes viruses subfamily [44], suggesting a unique function.

### The C-terminal SIM is critical to SUMO Targeted Ub Ligase activity of ORF61

Given that ORF61 includes both the RING domain and SIMs, we wanted to investigate STUbL activity of ORF61. We selected linear tetra SUMO2 as the substrate for the STUbL assay. The human STUbL RNF4 (aka SNURF) was used as a reference (positive control) [48]. In the polyubiquitination reaction, we added linear tetra-SUMO2 as the substrate. Ubiquitinated tetra-SUMO2 was synthesized by both ORF61 and RNF4, suggesting ORF61 binds to SUMO and assembles ubiquitin chains on them (Fig 6a). However, its activity was lower than RNF4. We performed the STUbL assay with the ORF61-RING mutants (Fig 6b). The STUbL activity of mutants correlated with their reduced polyubiquitination activity, highlighting the importance of the E2:RING contacts for STUbL activity.

**Figure 6:**
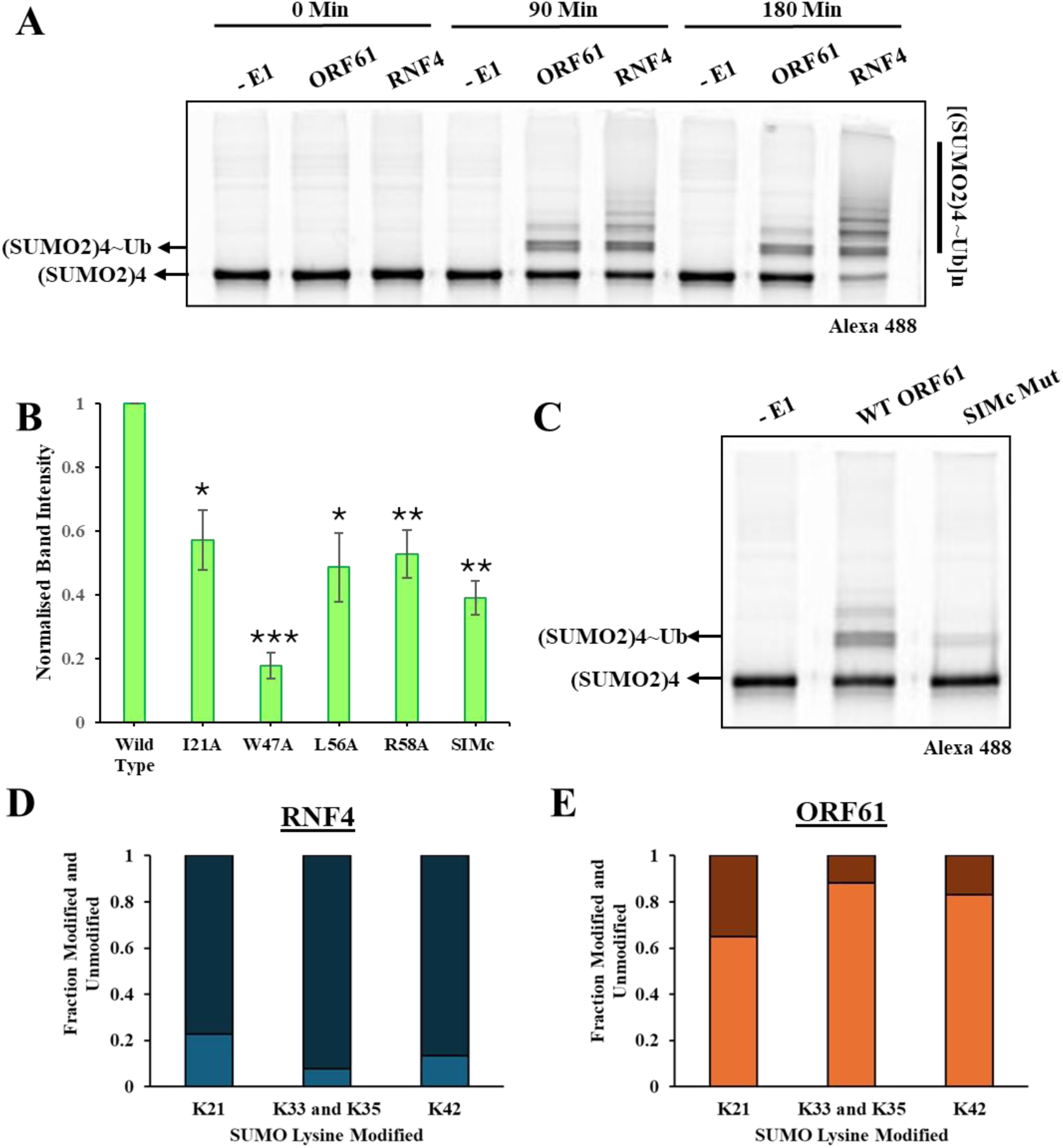
Assessment of STUbL Activity of ORF61 *in vitro*. A) We observed emergence of ubiquitinated species of tetra-SUMO2 over three hours suggestive of the SUMO Targeted Ubiquitin Ligase (STUbL) activity of purified ORF61 and RNF4 (positive control). B) Compared to the wild type ORF61, the emergence of mono-ubiquitinated tetra-SUMO2 ((SUMO2)4∼Ub) was found to be significantly reduced for all ORF61 mutants. For the RING domain mutants, the reduction was corresponding to the loss in activity observed for the respective E3 ligase activity. These assays were done in triplicates. Significant loss of ∼ 60% takes place in the STUbL activity of ORF61 takes place when the C-terminal SIM is mutated to all alanine and the SUMO - SIM hydrophobic interactions are disrupted as shown in C). D) and E) Comparative analysis of lysine ubiquitination preference between RNF4 and ORF61 *in vitro* at the end point of STUbL assay. The fraction of ubiquitin modified (-GG carrying) lysine peptides observed post

**Figure 07:**
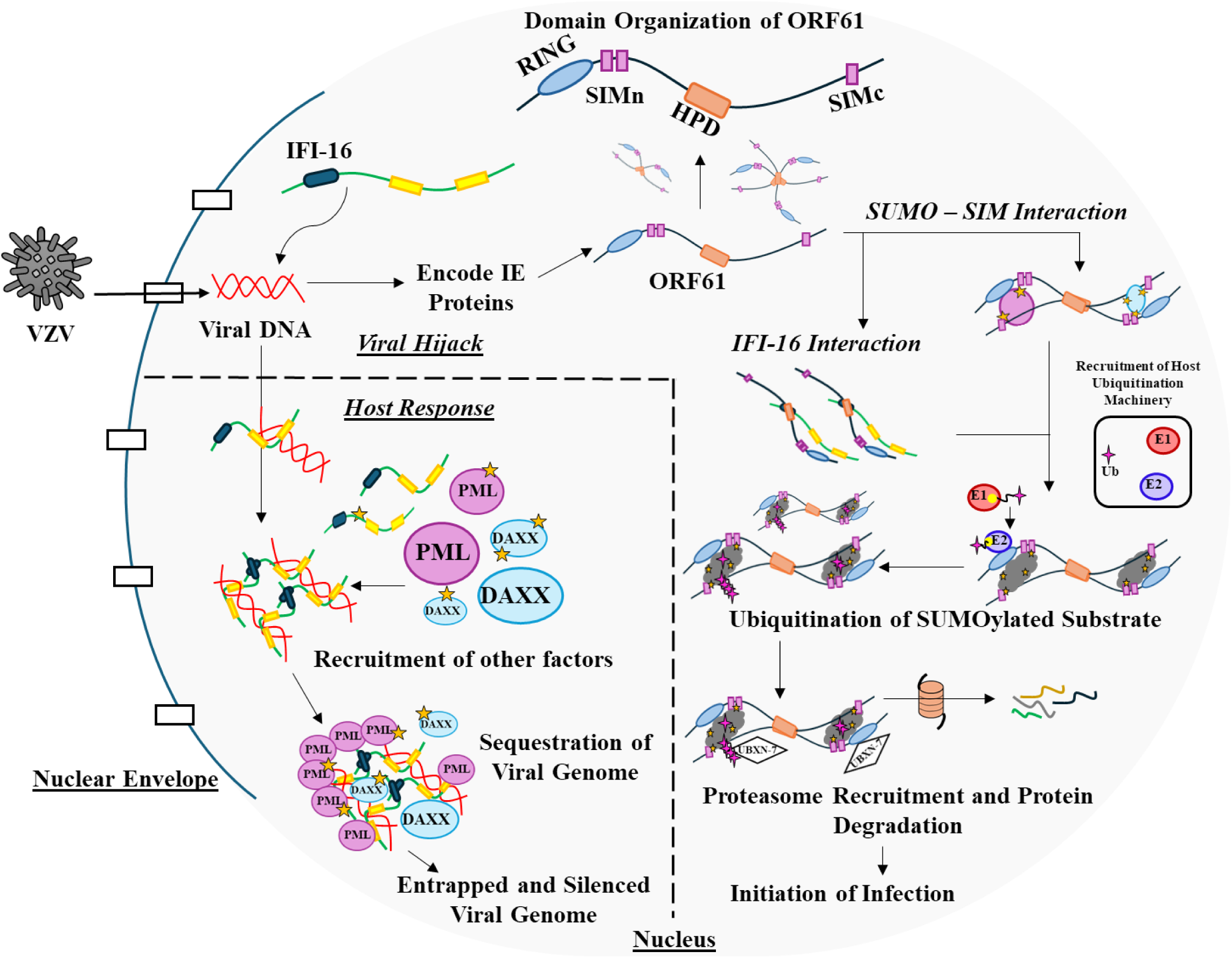
Proposed Mechanism of Action by ORF61 in the Host Cells. The known host response to exogenous genome entering the nucleus is the recruitment of IFI-16, which in this case sense the incoming viral genome of VZV and gets recruited to its site of entry. It interacts with the viral genome through two HIN200 (yellow boxes) while the Pyrin domain (PYD, dark blue box) facilitates its oligomerization. This response further elicits recruitment of other host innate immune response factors like PML and DAXX which lead to formation of newer PML nuclear bodies (PML-NBs) which ultimately sequester and silence the viral genome. Enrichment of SUMOylated proteins is a feature of PML-NBs depicted with yellow star onto PML and DAXX. ORF61 is one of the immediate early proteins encoded by the VZV genome. It carries an N-terminal RING domain and three SIMs with a central hydrophobic domain (HPD) annotated in the figure. The HDP potentially mediates self-oligomerization of ORF61 into dimers or higher order assemblies. The HPD of ORF61 could also interact with IFI-16 through PYD and sequester it to prevent its interaction with viral genome and/ or interact with SUMOylated forms of PML and DAXX and other components of PML-NBs through its SIMs and sequester them as the SIMc exhibits stronger affinity. This results in SUMOylated substrates (black cloud with yellow stars) in complex with ORF61 which subsequently recruits host ubiquitination machinery with the N-terminal RING domain. Being a ubiquitin E3 ligase, it is able to ubiquitinate the sequestered proteins (SUMOylated IFI16, PML, DAXX and other host response factors). This macromolecular complex then recruits the adaptor UBXN7, which recognizes the ubiquitin chains of the complex through its UIM motif or the ORF61 domain interacts directly with one of the other motifs of UBXN7. VCP/p97 and proteosome (orange barrel) ultimately degrade the sequestered protein complex allowing the viral genome to initiate infection.

To investigate the relevance of the C-terminal SIM for STUbL activity, we generated a SIMc mutant via site-directed mutagenesis in which the four hydrophobic residues were mutated to an alanine stretch. Compared with the wild-type ORF61, we observe a significant reduction in STUbL activity in the SIMc mutant (Fig. 6c). The loss in activity is almost 60% (Fig 6b), implying that SIMc contributes significantly towards ORF61’s STUbL activity. We further examined the specific target lysine residues of tetra-SUMO2, that are ubiquitinated by ORF61 and RNF4. using the Ub-clipping method discussed in the previous section. RNF4 modified K21, K33/K35, and K42 (Fig 6d). K33 and K35 are on the same peptide and could not be distinguished. ORF61 also ubiquitinated the same sites, albeit the extent of modification differed due to inherent differences in the STUbL activity (Fig 6e). Overall, SIMc is crucial for the STUbL activity of ORF61.

## Discussion

Pathogens have co-opted, mimicked, and disrupted the components of the host machinery to bypass innate immune response. In this study, we investigated how the immediate-early protein ORF61 encoded by Varicella-zoster virus (VZV) engages the host ubiquitin and SUMO signalling network. ORF61 is a RING-type ubiquitin E3 ligase [49], [50]. Our investigation reveals that ORF61 preferably interacts with the host E2D family of E2 enzymes (Fig 1). Despite the limited sequence conservation observed among the RING domain, ORF61 retains the classical β2α RING fold and engages with E2 through a canonical RING–E2 interaction network (Fig 2). The ORF61 RING critical network of residues at the E2:RING:Ub interface contributes to the ubiquitination activity (Fig 1 and 3), suggesting that ORF61 mimics the structural and functional mechanism of host RING ligases.

A notable feature of ORF61 revealed by our study is its capacity to assemble multiple ubiquitin chain types with a measurable fraction of branched architecture (Fig 4). The capacity of generating chains of varying architecture has been increasingly recognized among pathogenic E3 ligases [51]. Different ubiquitin chain topologies and linkages produce distinct signalling outcomes, and the ability of ORF61 to generate diverse ubiquitin linkages therefore suggests that the enzyme acts as a broad regulator of host signalling cascades during infection.

Our data has further revealed the importance of SIMs in guiding ORF61 STUbL activity. From our titration studies, the C-terminal SIM (SIMc) appears to play a dominant functional role due to its strongest binding affinity among the three SIMs of ORF61 (Fig 5), as the disruption of this particular motif significantly attenuated the STUbL activity (Fig 6c), suggestive of its potential role in driving substrate interaction. The SIM architecture of ORF61 is dispersed, in contrast to the tandem SIM architecture of host STUbL RNF4. This key difference is likely to limit avidity-driven SUMOylated substrate interaction and subsequent ubiquitination by ORF61, which helps explain the difference in activity observed (Fig 6a).

IFI-16 plays a central role in detecting foreign DNA and initiating the host’s innate immune response. VZV genome is sequestered by IFI-16 in the nucleus, which binds to dsDNA via HIN200 domain and subsequently recruits PML and DAXX, leading to de novo PML-NBs synthesis for viral genome silencing. Recent reports suggested that ORF61 regulates host IFI-16 levels, and additionally colocalises with UBXN7 [52]. IFI-16 is SUMOylated in the nucleus [53]. ORF61 could interact with SUMOylated IFI-16 via the HPD, which putatively interacts with IFI-16 Pyrin domain, and the SIMc domain which interacts with SUMO. Due to ORF61’s STUbL activity, IFI-16 will be ubiquitinated. Interaction of ORF61 with UBXN7 could facilitate recruitment of the AAA-ATPase p97 to ubiquitinated IFI-16, ensuring IFI-16 degradation by the p97-proteasome pathway. This mechanism could prevent the de novo synthesis of PML-NBs, providing a plausible explanation for the long-standing observation that ORF61 disperses PML nuclear bodies without directly degrading their components. This model suggests that while the molecular details differ between ICP0 and ORF61, the broader approach of overriding nuclear antiviral defences using immediate-early proteins as STUbLs is conserved among alpha herpesviruses.

We are scratching the surface regarding how viral STUbLs co-opt the host ubiquitin and SUMO pathway. The molecular details of the interaction between ORF61 and its substrates, and the ubiquitin linkages assembled on them by ORF61, and the subsequent fate of these substrates require further investigation. Uncovering ORF61’s molecular mechanism emphasizes the sophisticated ways in which herpesviruses manipulate host post-translational modification systems to promote viral replication.

## Acknowledgments

This work was supported by the Tata Institute of Fundamental Research, Department of Atomic Energy, Government of India, under project identification number RTI 4006. The NMR data were acquired at the NCBS-TIFR NMR Facility and Mass Spectrometry Facility, which are supported by the Department of Atomic Energy, Government of India, under project number RTI 4006.

