## Supplementary Information for "Investigating the activity of Varicella Zoster Virus (VZV) SUMO-targeted Ubiquitin Ligase ORF61"

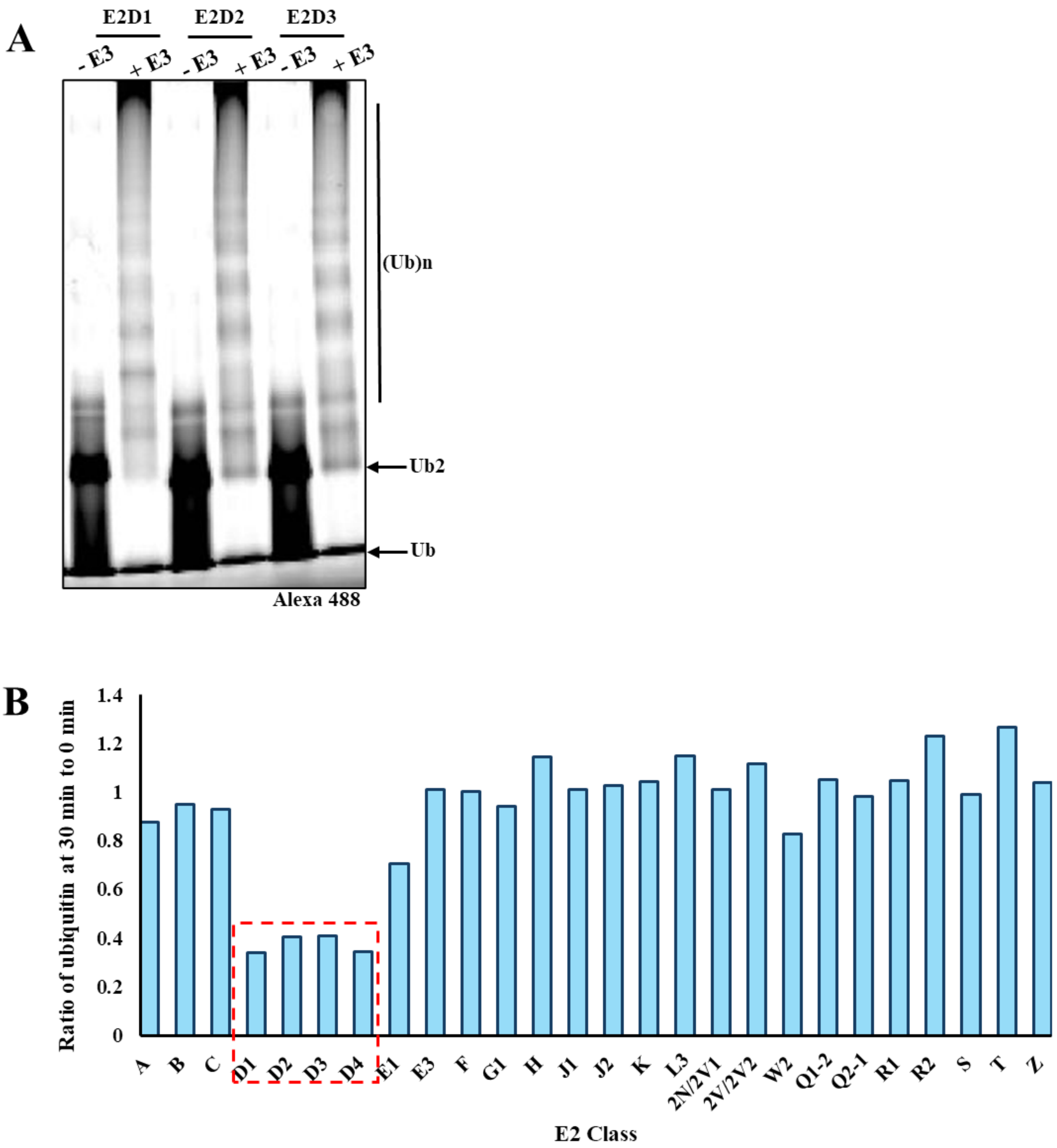


**Figure 1S:** Investigating activity of ORF61 with host E2 Enzymes.

A) High contrast image for E2D1, E2D2 and E2D3 from the E2 screen gel to emphasize the emergence of free polyubiquitin chains over 30 minutes. B) The plot shows the ratio of ubiquitin consumed in presence of E3 by comparing the ubiquitin band intensity for E2s from the screening performed. Here the reduced value is indicative of ubiquitin consumption as we are comparing the ratio of band intensity at 30 minutes to that of 0 minutes, lower value implies reduced amount of ubiquitin present at the end of 30 minutes suggestive of consumption during the assay. Minor sample loading error resulted in values more than 1 for E2H, E2L, E2 2V E2R2 and E2T classes.

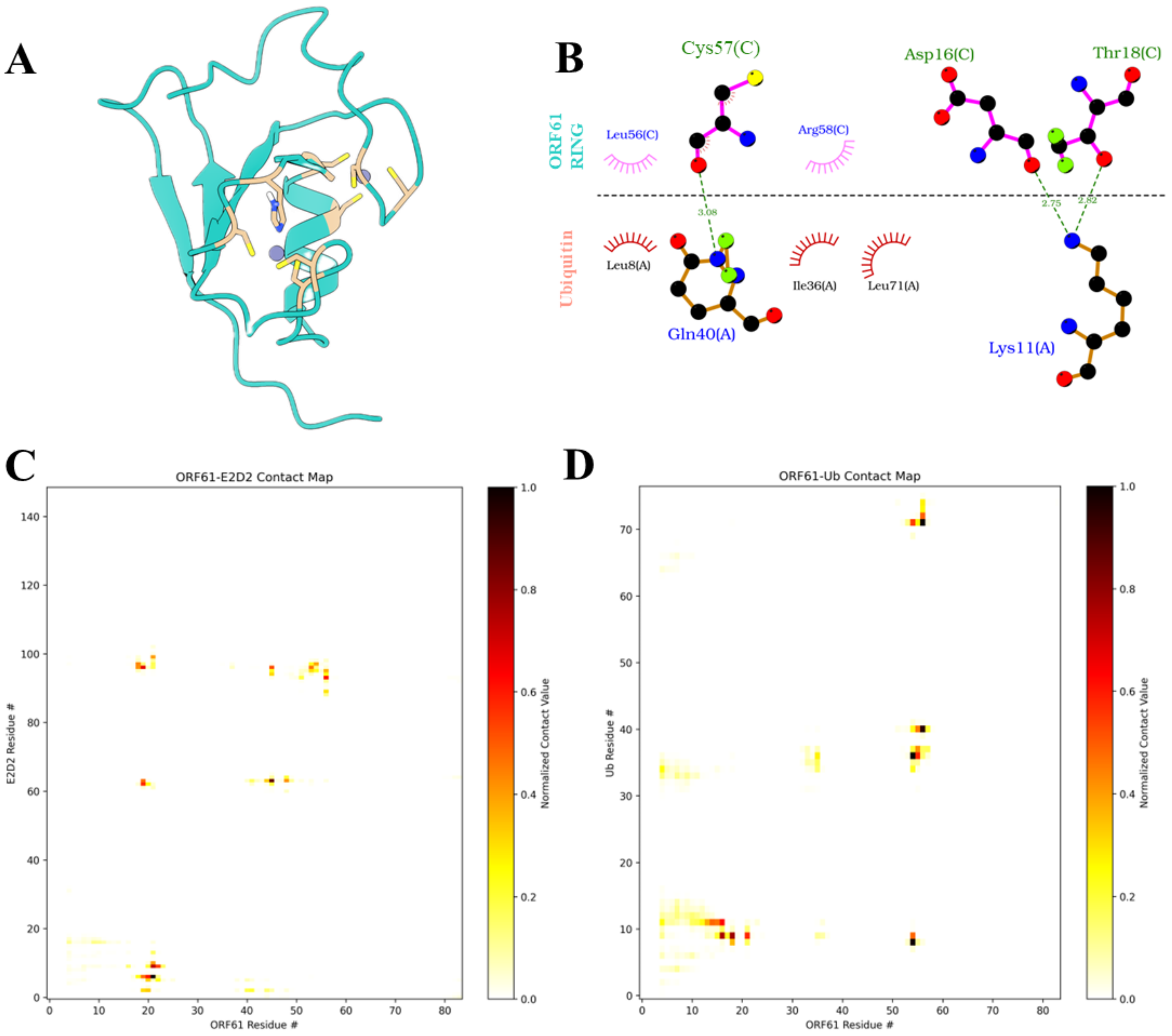


**Figure 2S:** Structure Modelling and Interface Analysis for ORF61 RING:E2~Ub Complex.

A) ORF61 RING domain of classic β2α fold in cross-brace zinc coordination scheme with zinc ions coloured in grey and residues coordinating to zinc ions are coloured in tan. B) The interaction map shows the residues of ORF61 RING and Ubiquitin interacting at the interface through hydrophobic (semi-circle representation) and hydrogen bond (dotted line representation) interaction. C) and D) Contact map of the residue interaction for ORF61 RING:E2D2 (C) and ORF61 RING:Ubiquitin (D). The representation is heatmap wherein the higher normalized score of contact value is indicated in darker colour which indicates higher prevalence of the two residues in contact throughout the simulation.

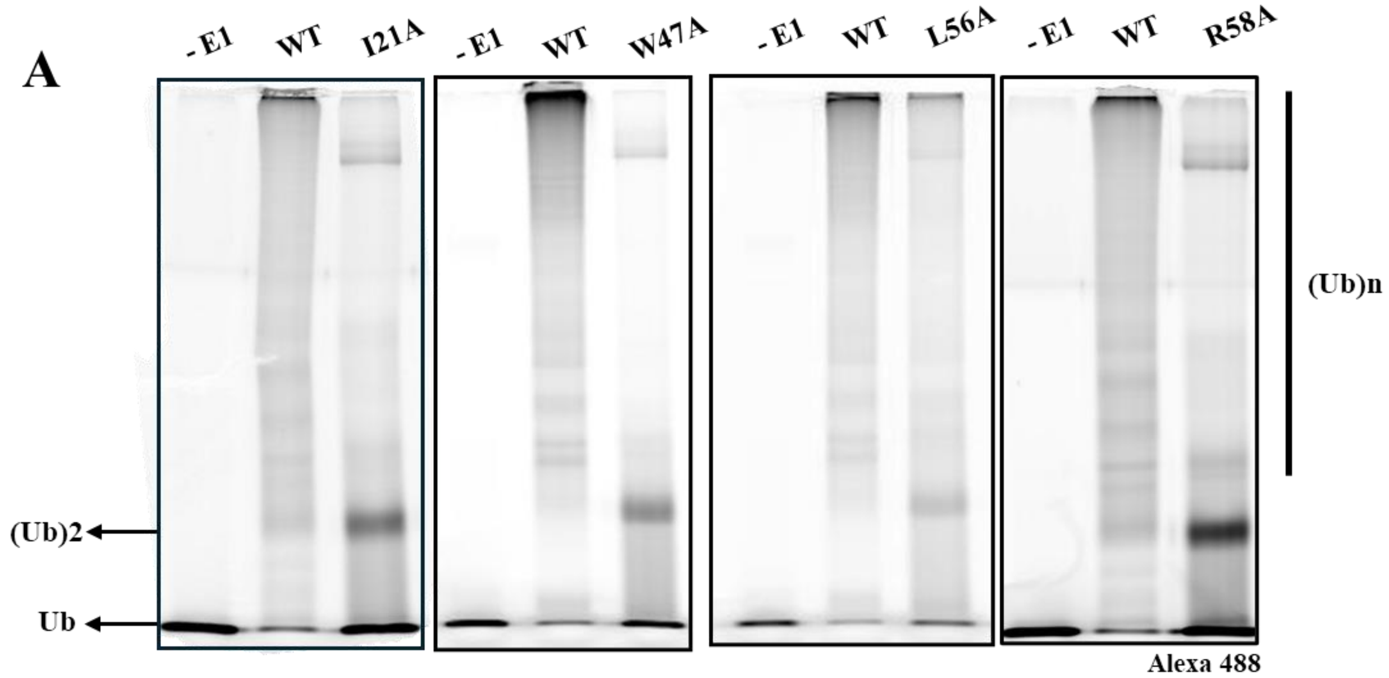


**Figure 3S:** Assessment of Functional Relevance of Residues at the RING - E2 Interface.

A) Polyubiquitination activity for all the RING mutants compared against wild type ORF61 at 30 minutes. No E1 was used as negative control.

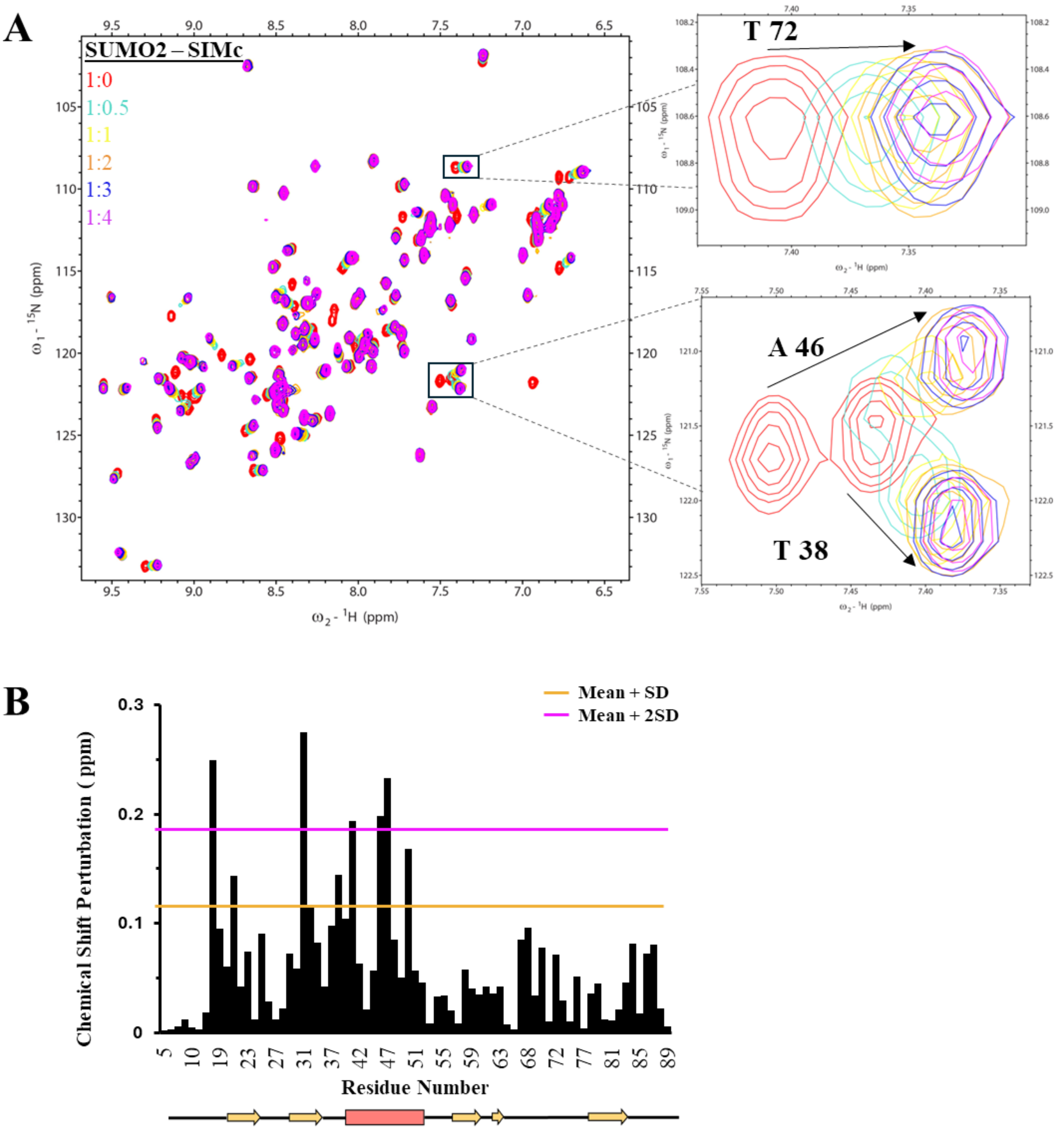

A
SUMO2 – SIMc
1:0
1:0.5
1:1
1:2
1:3
1:4
ω1 - 15N (ppm)
ω2 - 1H (ppm)
T 72
A 46
T 38
B
Mean + SD
Mean + 2SD
Chemical Shift Perturbation ( ppm)
Residue Number

C

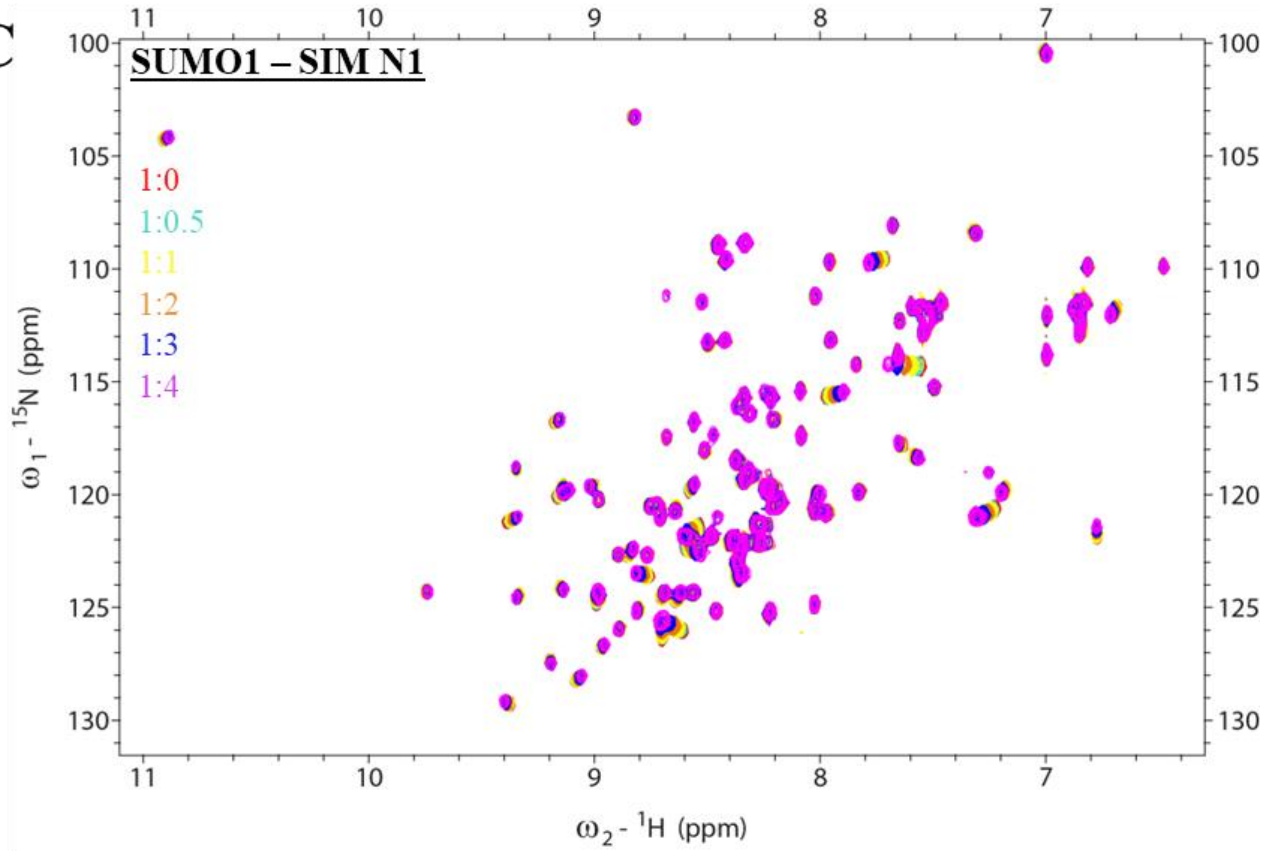

SUMO1 – SIM N1
1:0
1:0.5
1:1
1:2
1:3
1:4
ω1 - 15N (ppm)
ω2 - 1H (ppm)


D

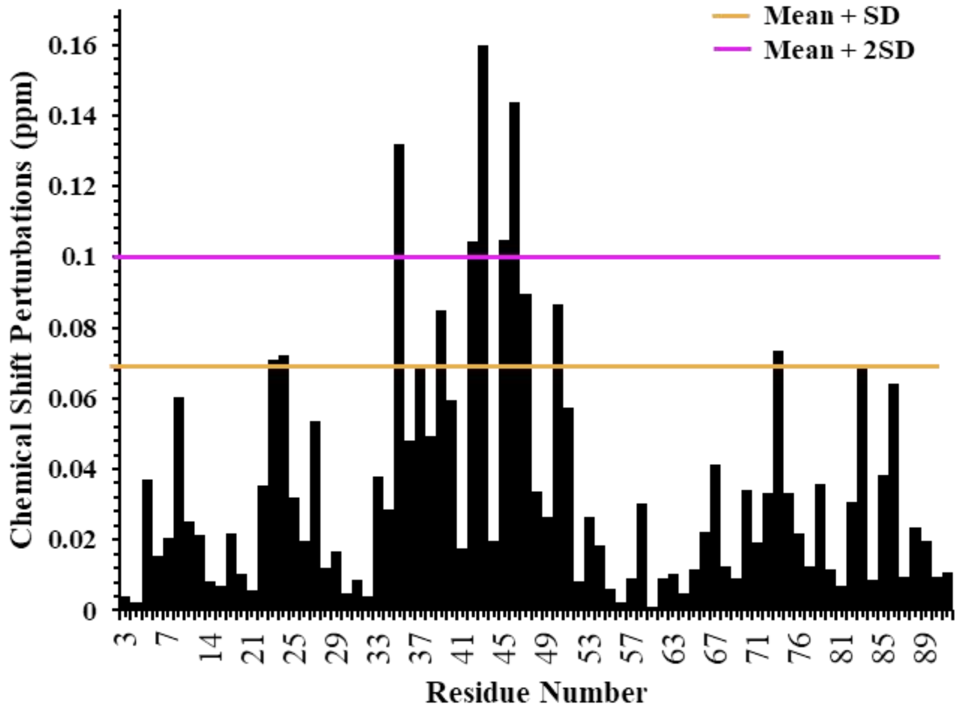

Mean + SD
Mean + 2SD
Chemical Shift Perturbations (ppm)
Residue Number

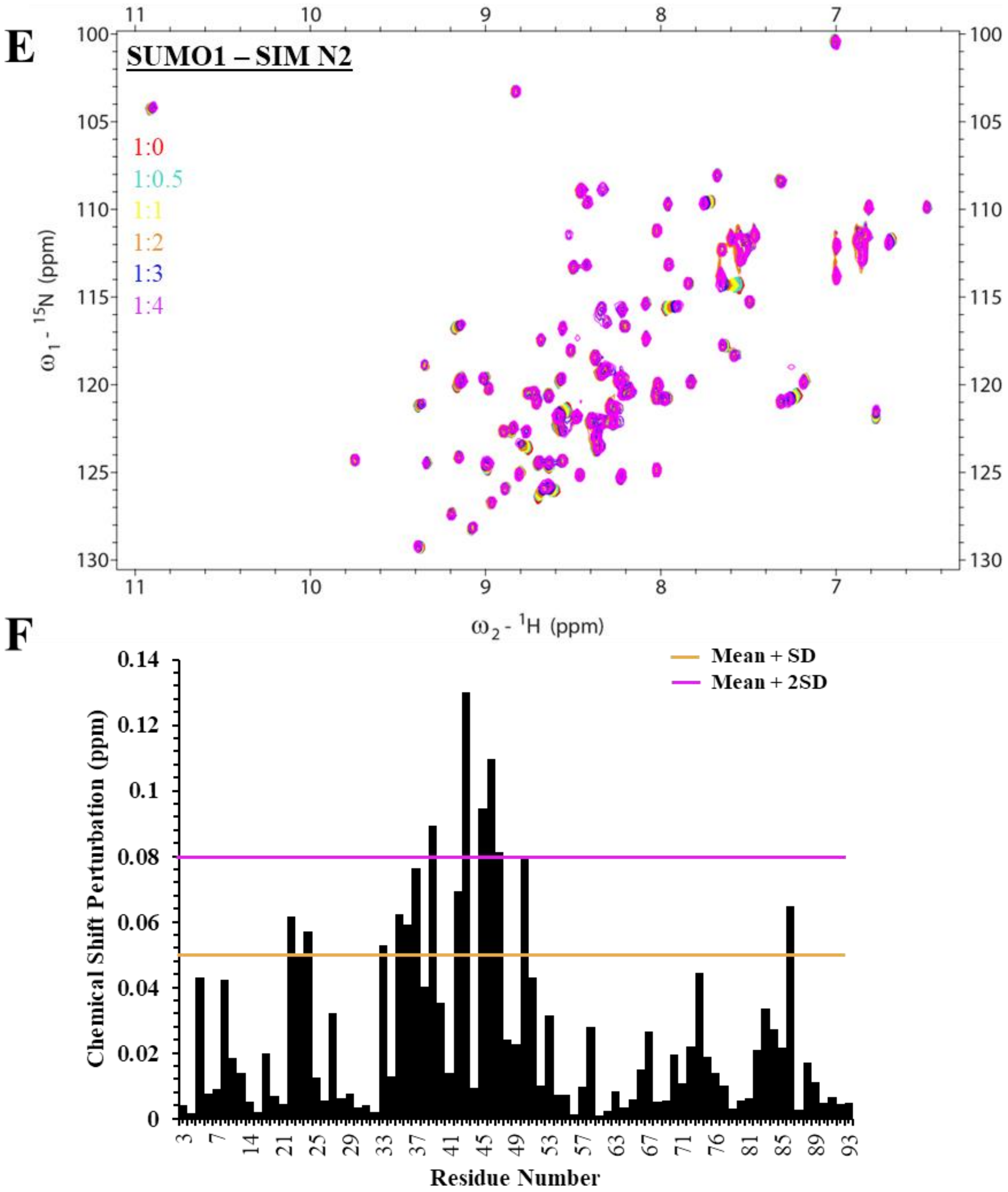
E
SUMO1 – SIM N2
1:0
1:0.5
1:1
1:2
1:3
1:4
ω1 - 15N (ppm)
ω2 - 1H (ppm)
F
Mean + SD
Mean + 2SD
Chemical Shift Perturbation (ppm)
Residue Number

G

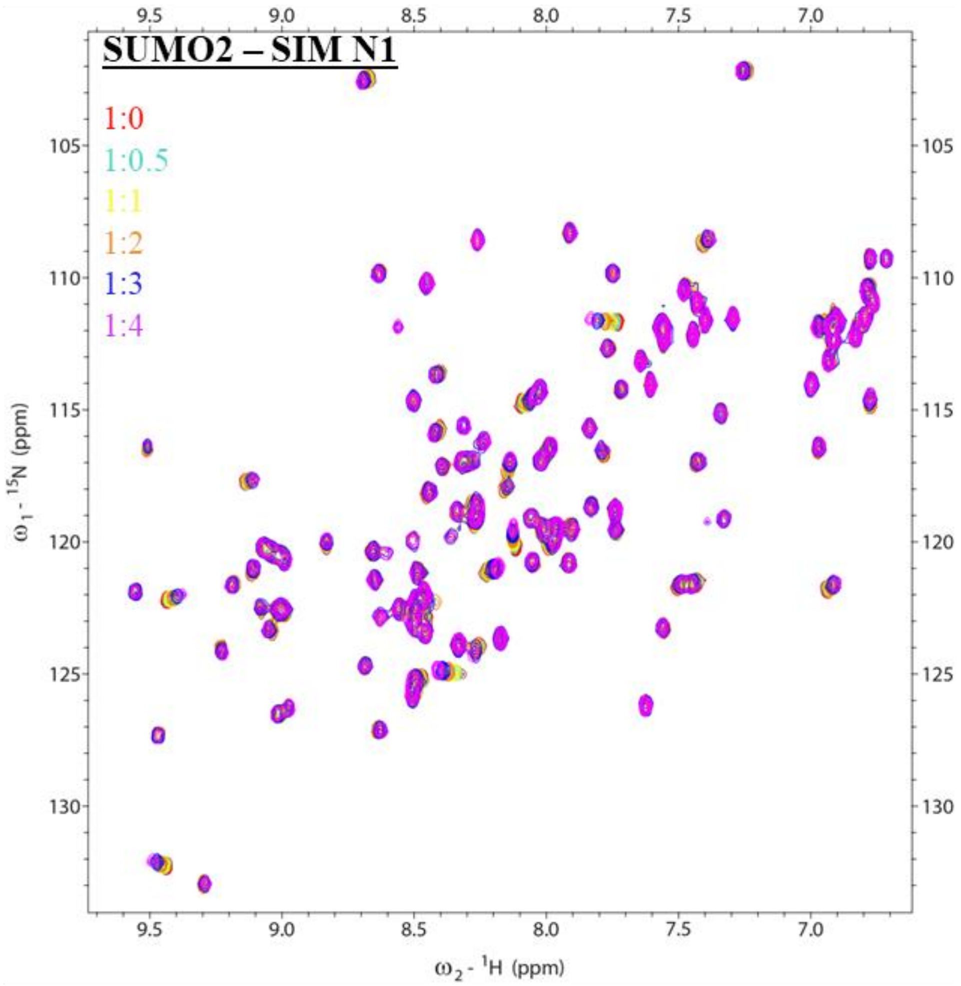
SUMO2 – SIM N1
1:0
1:0.5
1:1
1:2
1:3
1:4
$\omega_1$ - $^{15}$N (ppm)
$\omega_2$ - $^1$H (ppm)

H

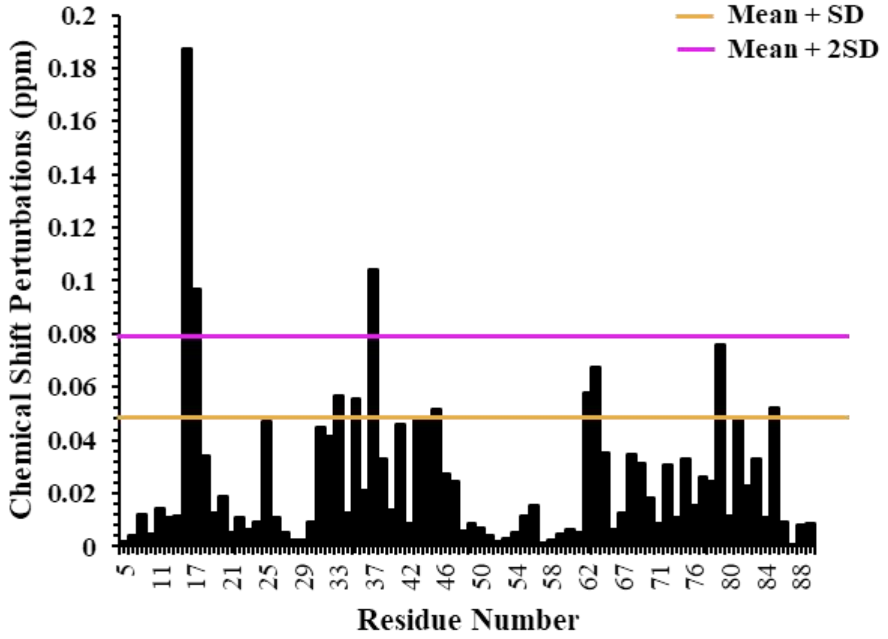
Mean + SD
Mean + 2SD
Chemical Shift Perturbations (ppm)
Residue Number

I

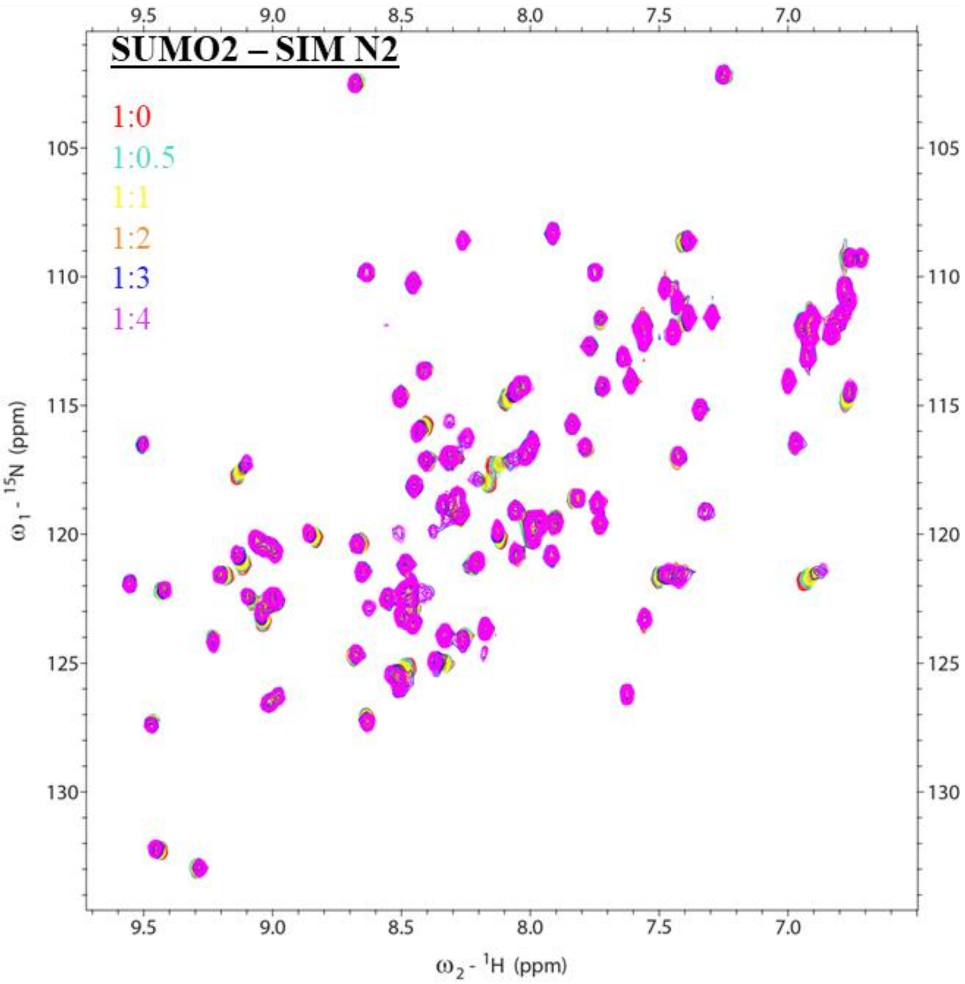
SUMO2 – SIM N2
1:0
1:0.5
1:1
1:2
1:3
1:4
ω1 - 15N (ppm)
ω2 - 1H (ppm)

J

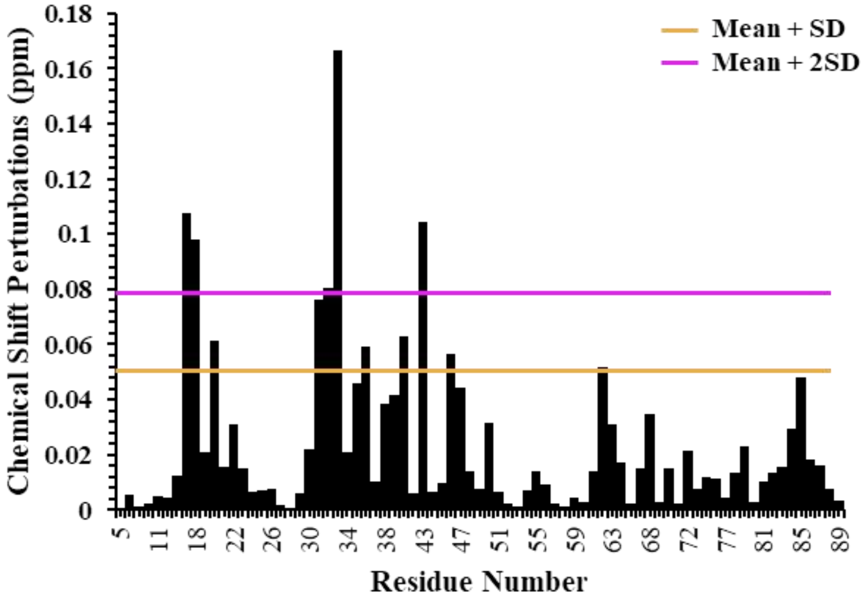
Mean + SD
Mean + 2SD
Chemical Shift Perturbations (ppm)
Residue Number

**Figure 5S:** Validating interaction of ORF61 SIMs with SUMO isoforms.

A) $^{15}$N-HSQC plot of SUMO2 isoform against increasing concentrations of ORF61 SUMO Interacting Motif (SIM) of C-terminal (SIMc) peptide at increasing concentrations. The inset shows two residues showing major perturbations. The chemical shift perturbation for all residues has been plotted in B). C) $^{15}$N-HSQC plot of SUMO1 isoform against increasing concentrations of the first SUMO Interacting Motif (SIM) of ORF61 at the N-terminal (SIM-N1) peptide at increasing concentrations. The inset shows two residues showing major perturbations. The chemical shift perturbation for all residues has been plotted in D). E) $^{15}$N-HSQC plot of SUMO1 isoform against increasing concentrations of the second SUMO Interacting Motif (SIM) of ORF61 at the N-terminal (SIM-N2) peptide at increasing concentrations. The inset shows two residues showing major perturbations. The chemical shift perturbation for all residues has been plotted in F). $^{15}$N-HSQC plot of SUMO2 isoform against increasing concentrations of the first SUMO Interacting Motif (SIM) of ORF61 at the N-terminal (SIM-N1) peptide G) and second SUMO Interacting Motif (SIM) of ORF61 at the N-terminal (SIM-N2) peptide I) at increasing concentrations. The inset shows two residues showing major perturbations. The chemical shift perturbation for all residues has been plotted in H) and J) respectively.
